# Spatial transcriptomics reveal markers of histopathological changes in Duchenne muscular dystrophy mouse models

**DOI:** 10.1101/2022.03.17.484699

**Authors:** L.G.M. Heezen, T. Abdelaal, M. van Putten, A. Aartsma-Rus, A. Mahfouz, P. Spitali

## Abstract

Duchenne muscular dystrophy (DMD) is caused by mutations in the *DMD* gene, leading to lack of dystrophin. Chronic muscle damage eventually leads to histological alterations in skeletal muscles. The identification of genes and cell types driving tissue remodeling is a key step to develop effective therapies. Here we use spatial transcriptomics in two DMD mouse models differing in disease severity to identify gene expression signatures underlying skeletal muscle pathologies and directly link this to the muscle histology. Deconvolution analysis allowed the identification of cell types contributing to histological alterations. We show how the expression of specific genes is enriched in areas of muscle regeneration (*Myl4*, *Sparc*, *Hspg2*), fibrosis (*Vim*, *Fn1*, *Thbs4*) and calcification (*Bgn*, *Ctsk*, *Spp1*). Finally, our analysis of differentiation dynamics in the severely affected *D2-mdx* muscle shows a subset of the muscle fibers are predicted to become affected in its future state. Genes associated with tissue remodeling could enable to design new diagnostic and therapeutic strategies for DMD.

## Introduction

Duchenne muscular dystrophy (DMD) is an X-linked recessive and fatal disorder affecting approximately one in 5000 male births ^1^. Out-of-frame mutations in the *DMD* gene on chromosome Xp21 lead to the absence of dystrophin protein ^2,3^. Dystrophin is part of the dystrophin-associated glycoprotein complex and acts as a linker between the extracellular matrix and the cytoskeleton, aiding in muscle membrane stability. The absence of dystrophin leads to a weakened sarcolemma, rendering the muscle fibers more vulnerable to contraction-induced damage ^4–7^.

Muscle fibers from early-stage DMD patients undergo cycles of degeneration and compensatory regeneration. Later, once the regenerative capacity of the muscle fibers becomes exhausted, muscle fibers are substituted by fibrotic and adipose tissue ^8–10^. Due to the accumulation of fibrotic and adipose tissue and subsequent progressive loss of muscle function ^11^, DMD patients become wheelchair dependent by 11-12 years of age and decease in their third or fourth decade due to cardiorespiratory failure ^12,13^.

Histological and molecular analyses of muscle samples obtained from DMD patients and animal models enabled the description of alterations resulting from lack of dystrophin. Studies in human showed how necrosis and inflammation promote the accumulation of fibrosis and fat infiltration ^11,14,15^. These observations were supported by bulk proteomics^16^ and transcriptomics studies^17^. Preclinical studies in the *mdx* mouse model (carrying a nonsense mutation in exon 23 of the *Dmd* gene) and especially in the more severely affected D2-*mdx* mouse (same mutation on a DBA/2J background), revealed the presence of necrosis, extensive inflammation, regeneration, central nucleation, changes in fiber size, fibrosis and calcified tissue ^18–20^. Various studies have assessed gene expression changes related to the disease. An upregulation of genes involved in inflammation (*Lgals3*, *Cd68*), muscle regeneration (*Myog*, *Pax7*, *Myh3*) and fibrosis (*Col1a1*, *Lox*) has been identified in preclinical models ^15,21–24^. These studies pointed out how specific cell types such as fibro-adipogenic progenitors (FAPs) and profibrotic factors (e.g. TGF-ß, CTGF and osteopontin) play an important role in regeneration and are thought to be main contributors of intramuscular fibrotic infiltration in dystrophic muscles ^25–30^.

Overall, associations between histological alterations with cell types and gene expression have been suggested. However, there are no studies where histological, cellular and gene expression data have been simultaneously and spatially investigated on the same tissue section. This limitation can be overcome by using spatial transcriptomics (ST), which allows direct linking of histology to gene expression. Here, we applied ST to identify molecular markers that underlie the histopathological changes observed in *mdx* and D2-*mdx* mice. We identified clusters based on gene expression profiling and histological features. Using spot deconvolution analysis, we show how different cell types contribute to the gene expression signature in tissue sections with 100 μm100μm resolution. Moreover, we identified differentially expressed marker genes that are involved in histopathological changes (regeneration, fibrosis and calcification). Finally, we predicted differentiation dynamics within the muscle section of the D2-*mdx* muscle using RNA velocity to assess whether we could identify a pattern predicting the future state of muscle fibers in tissue that is heavily affected by tissue remodeling^31^. Our study directly maps molecular changes to tissue alterations in preclinical models of DMD at spatial resolution the mapping of these data deepen our knowledge of DMD pathology, which facilitates development of future therapies for muscular dystrophies.

## Materials and method

### Animals

Mice were handled according to the guidelines established by the Animal Experiment Committee (Dierexperimenten commissie) of the Leiden University Medical Center (protocol PE.17.246.026). At 10 weeks of age, male C57BL/10ScSn-*Dmd*^mdx^/J (*mdx*),its matched wildtype background C57BL/10ScSnJ (C57BL10), the more severely affected mouse model *D2*.B10-*Dmd*^mdx^/J (D2-*mdx*) and its matched healthy background DBA/2J (DBA/2J) were euthanized by cervical dislocation.). A total of four mice were included in the protocol of which one mouse was used for Visium analysis. Mice were euthanized by cervical dislocation. The quadriceps (QUA) were isolated, embedded in O.C.T. compound (Sakura Finetek USA, Torrance, CA, USA), mounted on a piece of cork and fresh frozen in liquid nitrogen cooled isopentane. Tissue samples were transferred on liquid nitrogen and stored at −80°C until further processing.

### Visium spatial gene expression library construction

One fresh frozen isolated QUA (right-sided) from all four mouse models (C57BL10, DBA/2J, *mdx* and D2-*mdx*) was cryo-sectioned at 10 μm thickness at −21°C using a CryoStar NX70 cryostat (Thermo Scientific, Waltham, MA, USA). For each mouse model, one cross-section was placed on one of the four 6.5 mm-squared capture areas of a pre-cooled Visium Spatial Gene Expression slide (product code: 1000187, PN: 2000233, 10x Genomics) and adhered to the slide by warming the backside of the slide. The Visium Spatial Gene Expression slide was processed according to the manufacturer’s protocols. The slide was transferred on dry-ice and briefly warmed to 37°C for 1 minute after which fixation in ice-cold methanol was completed for 30 minutes at −20°C. Hereafter, the slide was covered with isopropanol at RT for 1 minute and consecutively a hematoxylin and eosin (H&E) staining (Agilent Technologies, Santa Clara, CA, USA; Sigma-Aldrich, Burlington, MA, USA) was performed according to the manufacturer’s protocol. Images covering the entire capture area were taken with a BZ-X700 microscope (Keyence, Osaka, Japan) using a 10× objective, stitched using the BZ-X700 analyzer software (Keyence) and exported as TIFF files. Permeabilization of the tissue sections was followed after imaging for 15 minutes. Upon release of the poly-adenylated mRNA from the tissue section, this was captured by barcoded primers on the Visium slide. Through reverse transcription in the presence of template-switching oligo, the captured mRNA was converted to spatially barcoded, full-length cDNA. Following second strand synthesis, a denaturation step released the cDNA from each capture area upon which PCR amplification, with a total of 16 cycles (based on Cq values) was performed. Subsequently, enzymatic fragmentation and size selection (SPRI beads) were used to optimize the cDNA amplicon size. Finally, to generate a sequencing ready indexed library, P5, P7, i7 and i5 sample indexes, and TruSeq Read 2 were added via end repair, A-tailing, adaptor ligation and PCR amplification. The spatial gene expression libraries were sequenced on an Illumina NovaSeq6000 with a target of ~125 million Paired-End reads per sample, or 50,000 reads per spot for each sample.

### Visium data processing and analysis

After sequencing, Illumina’s raw data was demultiplexed to fastq files using Space Ranger’s pipeline ‘mkfastq’ (10x Genomics Space Ranger v1.1.0). We used manual alignment of the tissue section and the fiducial frame (microscopic image) using the Loupe Browser (10x Genomics Loupe Browser v4.1.0).), followed by generating the spatial feature count matrices using Space Ranger’s pipeline ‘count’. Further analysis was done in R (version 4.1.1), using the Seurat package (version 4.0.5). Mitochondrial genes were filtered out before samples were further processed to overcome bias coming from these genes. The following genes were excluded: *mt-Nd1*, *mt-Nd2*, *mt-Co1*, *mt-Co2*, *mt-Atp8*, *mt-Atp6*, *mt-Co3*, *mt-Nd3*, *mt-Nd4l*, *mt-Nd4*, *mt-Nd5*, *mt-Nd6*, *mt-Cytb*. Furthermore, spots with very few counts or extremely high counts were excluded by filtering each sample separately based on the number of UMIs (nCount) and the number of genes (nFeature). The cut-offs were determined by visual inspection of the violin plots for each sample: for C57BL10 nCount >= 150 and <= 40000, nFeature >=150 and <=5000, for DBA/2J nCount >= 100 and <= 20000, nFeature >=200 and <=4000, for *mdx* nCount >= 200 and <= 40000, nFeature >=250 and <=5000 and for D2-*mdx* nCount >= 100 and <= 20000, nFeature >=150 and <=5000. Upon filtering, the spatial datasets were normalized using SCTransform. Hereafter, Principal Component Analysis was applied to the normalized data (RunPCA), and the top 30 principal components were used to generate a neighborhood graph using 20 neighbors (FindNeighbors). Next, data was clustered (FindClusters) with different resolutions (C57BL10 = 0.4, DBA/2J = 0.8, *mdx* = 0.8, D2-*mdx* = 0.4) and UMAP^32^ embedding was generated (RunUMAP). The FindAllMarkers (min.pct = 0.1, only.pos = FALSE) function was used to identify cluster-specific marker genes. The top 20 markers per cluster genes, based on Bonferroni-corrected p-values, were exported to EnrichR. Here, we looked at enrichment in cell types in the PanglaoDB Augmented 2021 reference dataset to guide the annotation of the clusters. The enriched cell types per cluster were used in the annotation of the clusters together with observation of the histological image underlying these cluster spots.

### Spot deconvolution using a single nucleus reference dataset

We used SPOTlight ^33^ to deconvolute each capture location (spot) of the 10X Visium data into cell types. SPOTlight uses seeded non-negative matrix factorization regression to integrate single-nuclei RNA sequencing (snRNAseq) and spatial transcriptomic datasets ^34^. SPOTlight learns topic signatures from a reference snRNAseq dataset and uses this to find the optimal weighted combinations of cell types to deconvolute the data from Visium spots into underlying cell types. We used the dataset of Chemello et al., 2020 as a reference snRNAseq dataset as this was the best matched single-nucleus dataset available ^35^. This snRNAseq dataset was obtained from the tibialis anterior (TA) of healthy wildtype (C57/BL6N) as well as an affected DMD mouse model (ΔEx51; deletion of *Dmd* exon 51). A total of *n* = 70 cells per cell type and *n* = 3000 highly variable genes were used as input for the deconvolution. Upon completion of the cell type deconvolution using SPOTlight, we calculated the average percentage of each cell type present in the previously annotated clusters. Moreover, we also calculated the average contribution of each cell type to the whole sample per mouse model.

### Identification of biomarkers using differential gene expression analysis

To identify biomarkers underlying histopathological changes in DMD mouse models, we performed differential gene expression analysis on a selection of spots. For each histopathological change of interest different marker genes and cut-offs were used to select these spots.

Regeneration was evaluated in the highly regenerative *mdx* mouse. Spots were considered to fall in the category ‘regeneration’ when they came from one of the two clusters: “regenerated fibers” or “regenerating fibers and inflamed patches” and were expressing either *Myog* and *Igfbp7* or *Myh3* and *Igfbp7* in the count data >0. The so called, non-regenerating category spots, came from the cluster “muscle fibers without CN” and did not express any of these markers. Fibrosis was evaluated in the more severely affected D2-*mdx* mouse. Spots were considered to fall in the ‘fibrosis’ category when they came from one of the following clusters: “inflamed and/or calcified fibers”, “muscle fibers” or “connective tissue” and were expressing either solely *Cd34*,or *Cd34* and *Lox* in the count data >0. Additionally, spots from the “connective tissue” cluster were included in the ‘fibrosis’ category when they expressed *Lox* and *Col1a1* to a higher extent with a threshold on the counts >3. The non-fibrotic fiber category was selected based on the “muscle fibers” cluster and no expression of *Cd34* or *Lox* and an expression <2 of *Col1a1*. Finally, calcification was also further investigated in the D2-*mdx* mouse. Spots were included in the ‘calcified’ category when they fell into the cluster “inflamed and/or calcified fibers” and had an expression >0 of the marker *Mgp*. The non-calcified category came from the cluster “muscle fibers” and had no expression of *Mgp*.

After selection of the categories, we assessed differentially expressed genes between the affected and unaffected spots using the FindMarkers function in Seurat. We used the default parameters in FindMarkers which uses a Wilcoxon Rank Sum test, only testing genes expressed in 10% of the cells in either group, and only testing genes with a log_2_(fold-change) > 0.25. Finally, we looked for positive (upregulated) markers in the affected category compared to the non-affected category. The genes were sorted based on fold-change and the top 50 marker genes were further evaluated.

### RNA velocity analysis

In order to apply the RNA velocity^31^ analysis, the unspliced and spliced gene expression are required for each spot. We used kallisto/BUStools^36,37^ to map the sequencing reads (fastq) to the reference genome (mm10) while quantifying intronic (unspliced) and exonic (spliced) reads. Next, we applied the RNA velocity pipeline^38^ (dynamic model implemented in the scvelo python package) to estimate the RNA velocity vector (i.e differentiation) of each spot in its spatial context. For the top 2000 variable genes, high dimensional RNA velocity vectors were calculated using 30 principal component and 30 neighbors. Next, these vectors were projected on the spatial coordinates of the tissue using the velocity_embedding and the velocity_embedding_stream functions.

To identify the genes mostly underlying the estimated RNA velocities, we considered the high dimensional RNA velocity vectors calculated using the top 2000 variable genes. For each spot, we transformed its high-dimensional RNA velocity vector into a unit vector (magnitude = 1) and we considered the squared value of each gene to represent the contribution of the gene to the RNA velocity of the spot of interest. For the ‘Inflamed and/or calcified fibers’ of the D2-*mdx* mice (324 spots), we calculated the top 5 contributing genes in each spot, and counted how many times each gene was observed in the top 5 list across all 324 spots.

## Results

### Spatial transcriptomics on skeletal muscle reveals clusters related to histology

Quadriceps muscle of healthy (C57BL10 and DBA/2J) and dystrophic (*mdx* and D2-*mdx*) mice were analyzed. The four muscles composing the quadriceps group (rectus femoris (RF), vastus lateralis (VL), vastus medialis (VM) and vastus intermedius (VI)) were present on the slide, although the VI was not present in all sections. Muscles of healthy mice showed homogenous tissue composition, with similarly sized muscle fibers, by H&E staining compared to dystrophic tissues, where variation in fiber size, necrosis, inflammation, fibrosis, regeneration (*mdx*) and calcification (D2-*mdx*) were observed (Figure 1A,D,G,J). After processing, a total of 474,186,322 paired-end reads covering 7028 spots were obtained with a median of 1122 genes per spot (Table 1).

**Figure 1.**
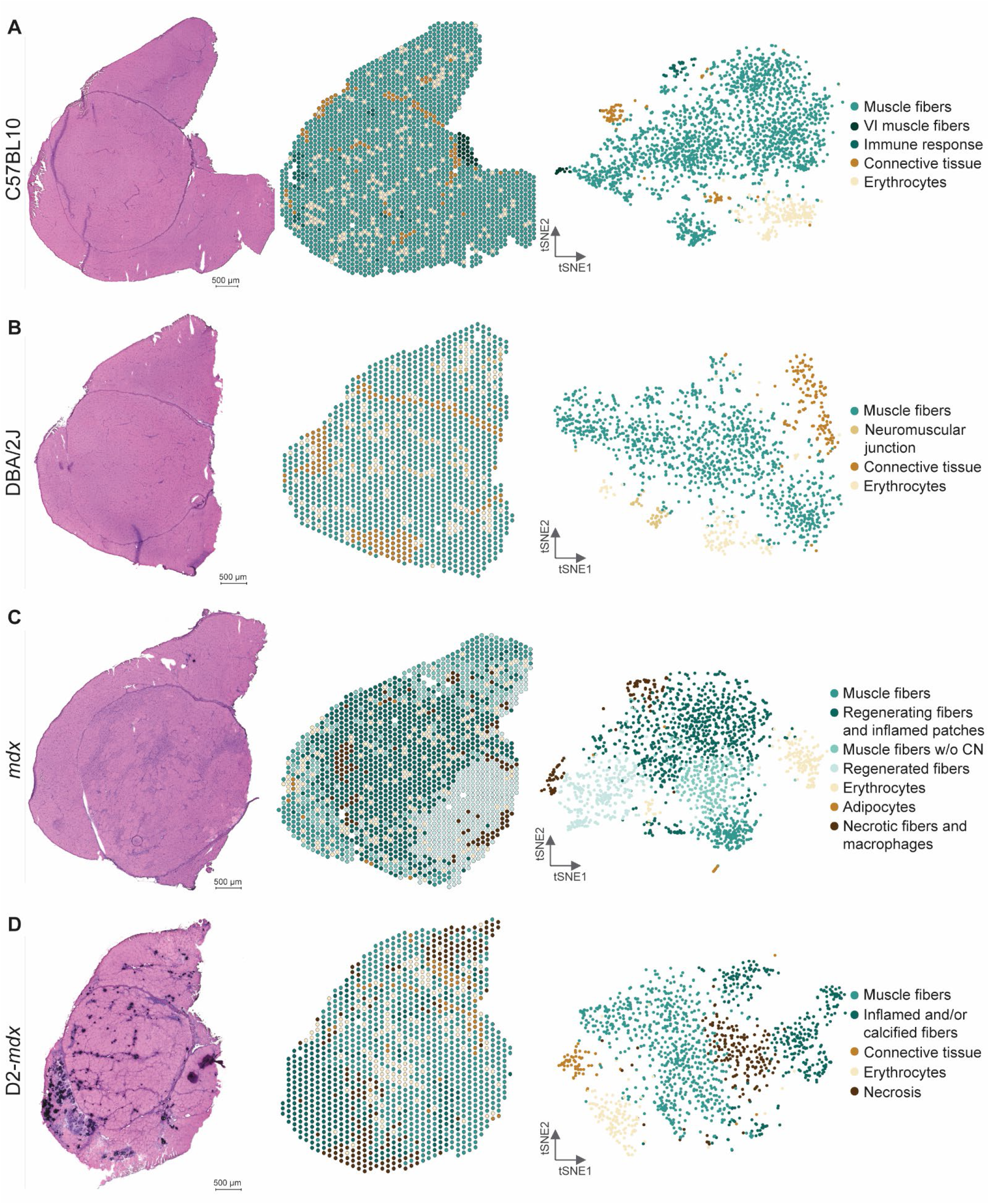
Characterizing muscle tissue in DMD and WT mouse models using spatial transcriptomics Characterizing muscle tissue in DMD and WT mouse models using spatial transcriptomics. (left) is the Hematoxylin and Eosin (HE) stained section, (middle) displays the Visium spots spatially plotted and colored by cluster, and (right) is a tSNE map of the Visium spots colored by cluster for (A) C57BL10, (B) DBA/2J, (C) mdx and (D) D2-mdx.

**Table 1.**
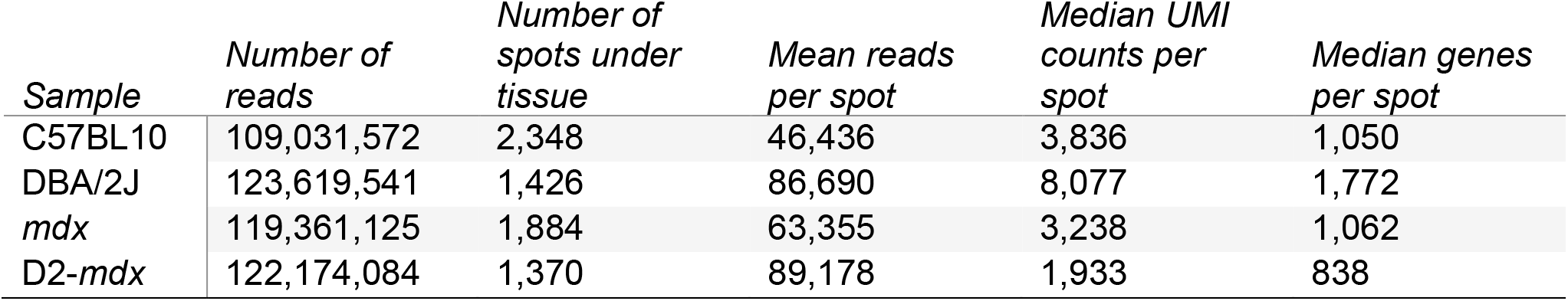
Sequencing results of Visium experiment.

| Sample | Number of reads | Number of spots under tissue | Mean reads per spot | Median UMI counts per spot | Median genes per spot |
| --- | --- | --- | --- | --- | --- |
| C57BL10 | 109,031,572 | 2,348 | 46,436 | 3,836 | 1,050 |
| DBA/2J | 123,619,541 | 1,426 | 86,690 | 8,077 | 1,772 |
| *mdx* | 119,361,125 | 1,884 | 63,355 | 3,238 | 1,062 |
| D2-*mdx* | 122,174,084 | 1,370 | 89,178 | 1,933 | 838 |

Spots were clustered for each sample independently, based on gene expression clusters were annotated based on marker genes and histological observations (see Methods for details). Clusters representing healthy muscle fibers were identified in the wildtype models. Top markers for healthy muscle fibers included *Myh4*, *Tpm2*, *Ckm*. We also identified clusters representing the perimysium and other connective tissue, expressing *Col1a2*, *Col1a1*, *Fmod*. A VI specific cluster was defined by the expression of myosins specific for slow twitch muscle fibers (*Myl2, Myl3, Myh7*) (Figure 1A, Supplementary Figure S1). Additionally, we identified smaller clusters enriched in genes specific for erythrocytes (*Hba-bt*, *Hbb-a2*, *Hba-a1*) and immune response (*Lyz2*, *Spp1*, *Ctsb*). The latter one was observed in an area of damaged tissue in the C57BL10 mouse only, underpinning the strength of using spatial transcriptomics to identify small tissue alterations (Supplementary Figure 1). An additional small cluster enriched for neuromuscular junction related genes (*Mpz*, *Mbp*, *Chrne*) was found in the DBA/2J muscle (Figure 1B, Supplementary Figure S2). Some clusters (muscle fibers, erythrocytes, connective tissue) were present in all mouse models with comparable top marker genes (Supplementary Table S1-4).

Dystrophic samples displayed additional clusters, not present in the control samples, matching the histopathological changes present in these samples. *Mdx* mice showed mature muscle fibers (*Amd1*, *Amd2*, *Smox*), regenerated fibers (*Gsn*, *Fhl1*, *Igfbp7*), regenerating fibers with inflamed patches (*Xirp2*, *Lars2*, *Mybpc1*) and some necrotic muscle fibers with infiltrating macrophages (*Cd68*, *Vim*, *Tyrobp*) (Figure 1C). Additionally, a small cluster of adipocytes was identified (*Apoc1*, *Scd1*, *Adipoq*) (Supplementary Figure S3). The more severely affected D2-*mdx* mouse additionally showed a large cluster with prominent inflammation and calcification signatures (*Spp1*, *Mgp*, *Mpeg1*) as well as necrotic fibers (*Uba52*, *Eef1d*, *Cox6a2*) (Figure 1D, Supplementary Figure S4).

### Unravelling the spot: enrichment of specific cell types in annotated clusters

The resolution of the Visium slides does not allow profiling gene expression at the single cell level. To overcome this and to estimate the contribution of different cell types to the observed transcriptomic signatures, we deconvoluted the spots using a snRNAseq reference dataset. This dataset was obtained from TA muscles of wildtype mice and a mouse model carrying a deletion of *Dmd* exon 51 (ΔEx51) resulting in dystrophin absence and histological alterations similar to those observed in *mdx* mice^35^. The selected reference dataset includes nuclei originating from muscle as well as non-muscle nuclei. These nuclei were annotated, allowing for deconvolution of several cell types (Figure 2A).

**Figure 2.**
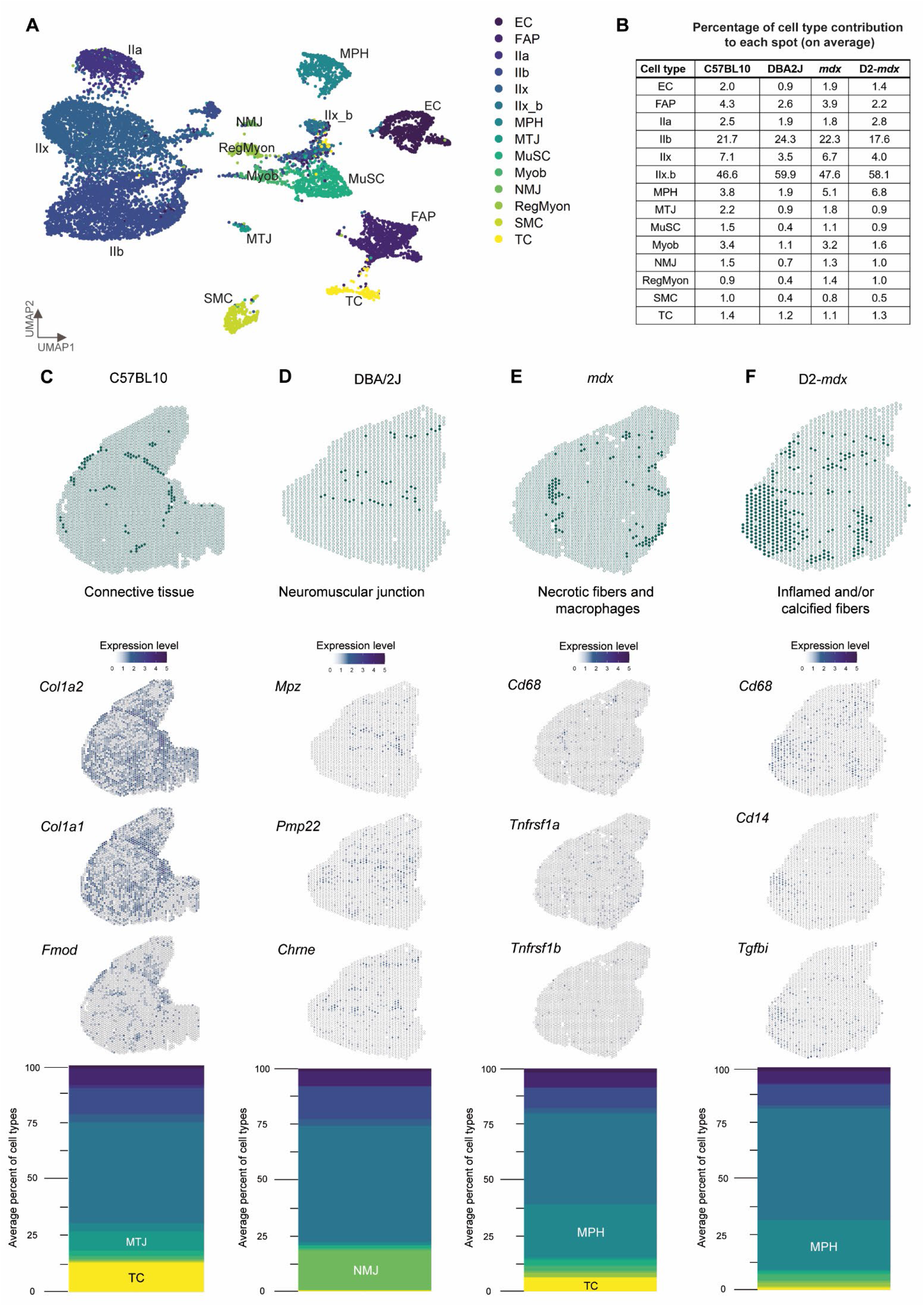
Deconvolution of the spatial data using a snRNAseq reference dataset reveals enrichment of cell types in different mouse models and specific clusters. (A) UMAP of reference dataset including the following cell types: endothelial cells (EC), fibro/adipogenic progenitors (FAP), type IIa myonuclei (IIa), type IIb myonuclei (IIb), type IIx myonuclei (IIx), type IIx_b myonuclei (IIx_b), macrophages (MPH), myotendinous junction myonuclei (MTJ), muscle satellite cells (MuSC), myoblasts (Myob), neuromuscular junction myonuclei (NMJ), regenerative myonuclei (RegMyon), smooth muscle cells (SMC) and tenocytes (TC) (B) The average percentage of cell type contribution per spot in the four mouse models (C) Deconvolution of the “connective tissue” cluster from C57BL10 showing marker genes Col1a2, Col1a1 and Fmod and a stacked barplot displaying the average percentage of contributing cell types to this cluster with an enrichment for MTJ and TC (D) Deconvolution of the “neuromuscular junction” cluster from DBA/2J plotted marker genes Mpz, Pmp22 and Chrne and a stacked barplot displaying the average percentage of contributing cell types to this cluster with an enrichment for NMJ (E) Deconvolution of the “necrotic fibers and macrophages” cluster from mdx, with marker genes Cd68, Tnfrs1a and Tnfrs1b and a stacked barplot displaying the average percentage of contributing cell types to this cluster with an enrichment for MPH and TC (F) Deconvolution of the “inflamed and/or calcified fibers” cluster from D2-mdx, with marker genes Cd68, Cd14 and Tgfbi and a stacked barplot displaying the average percentage of contributing cell types to this cluster with an enrichment for MPH.

Deconvolution showed how the different cell types contribute consistently to the gene expression signature. The main contributing cell types were type IIx myonuclei (ranging from 46.6% to 59.9% of nuclei across the 4 quadriceps muscles, Figure 2B), followed by type IIb myonuclei (ranging from 17.6% to 24.3% of nuclei). A consistent increase in macrophages (MPH) was observed in *mdx* and D2-*mdx* mice compared to their wildtypes. Furthermore, the muscle regenerative capacity characteristic of the BL10 background was reflected by elevated levels of myoblasts, muscle satellite cells and regenerative myonuclei compared to the mice on the DBA/2J background known to have reduced regeneration potential.

The average cell type contribution to the spots assigned to each specific cluster was calculated to assess whether there is an enrichment for specific cell types across annotated clusters. Figure 2C shows how the connective tissue cluster in the C57BL10 muscle matches the distribution of markers (*Col1a2*, *Col1a1* and *Fmod*) associated with the connective tissue, tenocytes (TC) and the myotendinous junction myonuclei (MTJ). Deconvolution showed an enrichment for these cell types at these locations, with 12.9% of nuclei being TC and 8.5% of nuclei being MTJ compared to an overall respective average of 1.4% and 2.2% per spot in the C57BL10 sample.

Another example is given by zooming intoto the neuromuscular junction (NMJ) cluster in the DBA/2J tissue section (Figure 2D). Mapping the specific marker genes for the NMJ myonuclei (*Mpz*, *Pmp22* and *Chrne*) shows increased expression in the cluster spots. This is also reflected upon deconvolution, where the NMJ myonuclei are a clear contributing factor for the spots included in these clusters. The average percentage of NMJ myonuclei per spot was 0.7% for the DBA/2J model, while being 17.8% in these selected spots.

Next, we focused on the clusters that showed clear histopathological changes in both the *mdx* and D2-*mdx* mice (necrotic fibers and macrophages, inflamed and/or calcified fibers) (Figure 2E-F). Macrophage markers (*Cd68* for M1-macrophages, *Cd14* for M2-macrophages and *Tgfbi* as a secretion marker of M2-macrophages) were plotted accompanied by necrosis specific markers (*Tnfrsf1a* and *Tnfrsf1b*). The expression pattern of these markers spatially aligned with the spots in these clusters. In *mdx* mice, an enrichment for MPH in these spots (23.8% of nuclei) was observed compared to the average per spot (5.1%). Finally, in D2-*mdx* the overall average per spot of MPH contribution was 6.8% and in the assigned cluster 22.3%. Altogether, deconvolution analysis reveals the cell types underlying the observed gene expression patterns in the Visium spots.

### Identification of muscle regeneration biomarkers

From 3 to 12 weeks of age, *mdx* muscles undergo repetitive cycles of degeneration and regeneration ^39^. To identify genes associated with muscle regeneration we compared areas characterized by the presence of centralized nuclei (CN) with areas with nuclei located at the periphery in the *mdx* mouse (Figure 3A). Spot selection was based on previously defined clusters (regenerating: regenerated fibers and regenerating fibers and inflamed patches, non-regenerating: muscle fibers without CN). An additional criteria for selection was the expression of *Myog*, *Igfbp7* and *Myh3* in areas of regeneration and absence of these genes in non-regenerating spots (Figure 3B). Differential gene expression analysis between these two groups of spots revealed markers of regeneration (top 50 markers summarized in Table S5) such as *Myl4*, *Sparc* and *Hspg2*, which showed little-to-no expression in the non-regenerating spots (Figure 3C). Spatially, the expression of these genes located in areas of regeneration as marked by the histology (Figure 3D). Besides upregulation of neonatal myosins (*Myl4*, log_2_(FC) = 1.42, adjusted p-value = 5.50e-17), we also saw upregulation of *Sparc* ^40,41^ only in areas marked as regenerating muscle fibers (log_2_(FC) = 0.90, adjusted p-value = 5.61e-16, Figure 3D-*mdx*). Finally, we identify *Hspg2*, a gene involved in cell growth and differentiation, as an important upregulated molecular marker in areas of regeneration (log_2_(FC) = 0.57, adjusted p-value = 1.76e-15). To evaluate the specificity of the association of the identified genes with regeneration we assessed the expression of these genes in C57BL10 wildtypes where muscle regeneration was low. While *Myl4* expression was very low in wildtype mice, *Sparc* and *Hspg2* expression was not negligible in wildtype mice. Especially *Sparc* expression was observed at high levels across the tissue, but primarily in the connective tissue sheet and VI muscle (Figure 3D-C57BL10).

**Figure 3.**
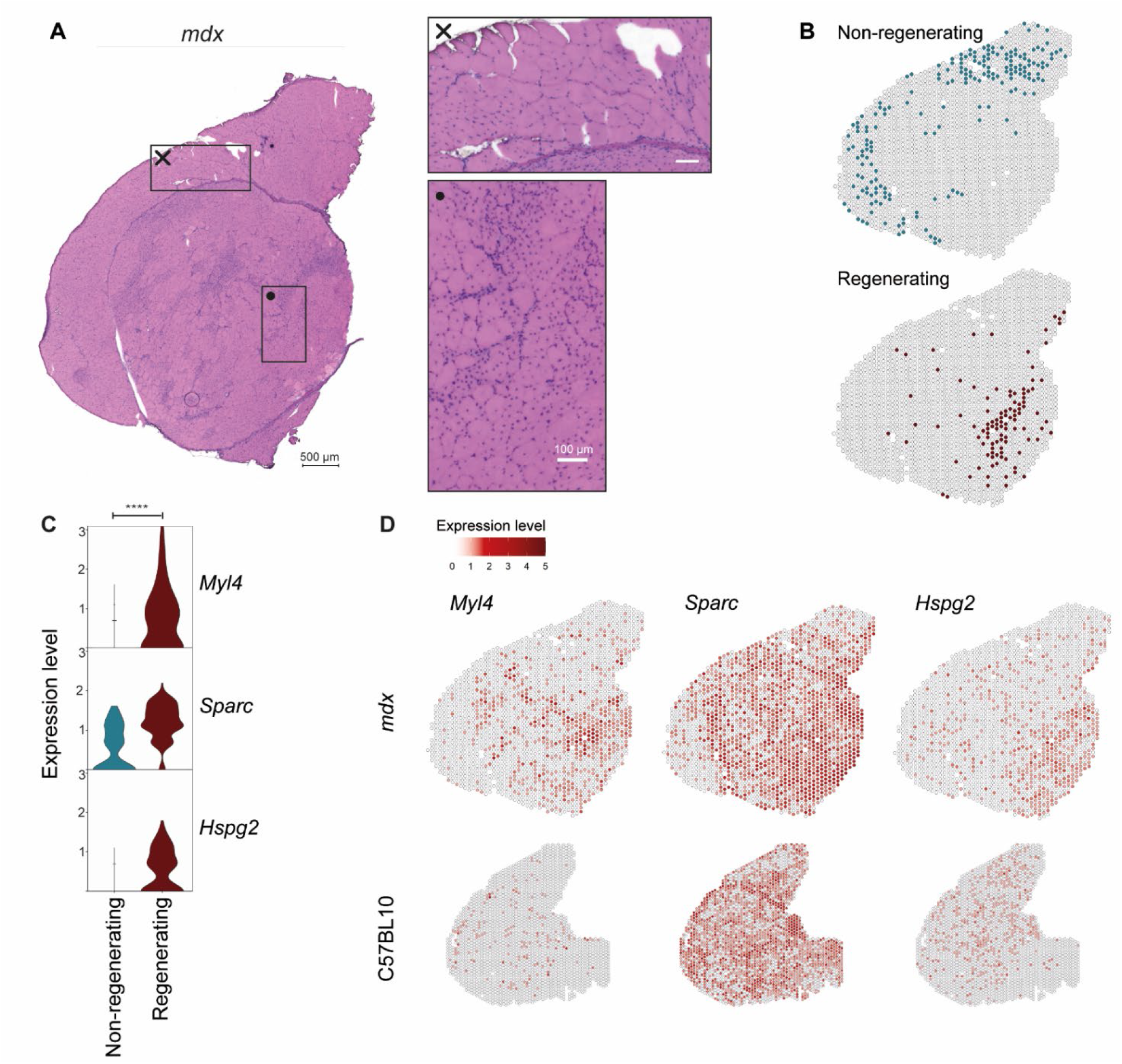
Identifying biomarkers of muscle regeneration in the mdx mouse model. (A) HE stained mdx QUA sample with two zoomed-in areas, × displaying an area of muscle fibers without centralized nuclei (CN) which are thought to be not regenerated yet, • showing an area of recently regenerated or regenerating muscle fibers (B) Selected spots belonging to the categories “non-regenerating” or “regenerating” were compared for differential gene expression analysis (C) Expression levels of selected genes Myl4, Sparc and Hspg2 in the two categories with an enrichment in the “regenerating” spots (**** representing a p-value <0.0001) (D) Spatially plotted expression of Myl4, Sparc and Hspg2 in mdx and C57BL10 muscle.

### Identification of genes underlying severe dystrophic histopathological changes

Given the limited severity of the *mdx* model, we sought to identify genes associated with fibrosis, changes to tissue remodeling and calcification in the D2-*mdx* model. To identify fibrotic markers, spots expressing known fibrotic genes such as *Cd34*, *Lox* and/or *Col1a1* were compared to spots belonging to the muscle fibers cluster where fibrotic genes were not expressed (Figure 4B). The top 50 differentially expressed genes are summarized in Table S6. Genes such as *Vim*, *Fn1* and *Thbs4* showed significant enrichment in fibrotic regions(Figure 4D). *Vim* was expressed in non-fibrotic spots, but at significantly lower levels compared to fibrotic spots (log_2_(FC) = 1.11, adjusted p-value = 1.71e-10). *Vim* encodes for the myofibroblastic protein vimentin and is expected to be upregulated in areas of skeletal muscle fibrosis ^42–44^. Given the role of *Vim* in focal adhesion, where regulatory signals and mechanical force is transmitted between the extracellular matrix (ECM) and an interacting cell ^45^, expression was found in healthy tissue areas as well (Figure 4E-DBA/2J). The next identified biomarker of fibrotic infiltration is *Fn1* encoding for fibronectin. Co-expression of *Fn1* and *Vim* is often used to identify fibroblasts and is thus a known marker involved in this histopathological change ^46,47^. *Fn1* was highly upregulated in fibrotic spots (log_2_(FC) = 1.38, adjusted p-value = 1.80e-16, Figure 4D) and spatially mapped the locations where the tissue damage was most severe (Figure 4E-D2-*mdx*). Finally, *Thbs4* seemed more specific for fibrosis rather than being present in all damaged areas, given the more restricted expression pattern in D2-*mdx* (log_2_(FC) = 1.32, adjusted p-value =1.46e-16). Whereas, localization of *Thbs4* expression is limited to perimysial tissue in the healthy DBA/2J sample (Figure 4E-DBA/2J).

**Figure 4.**
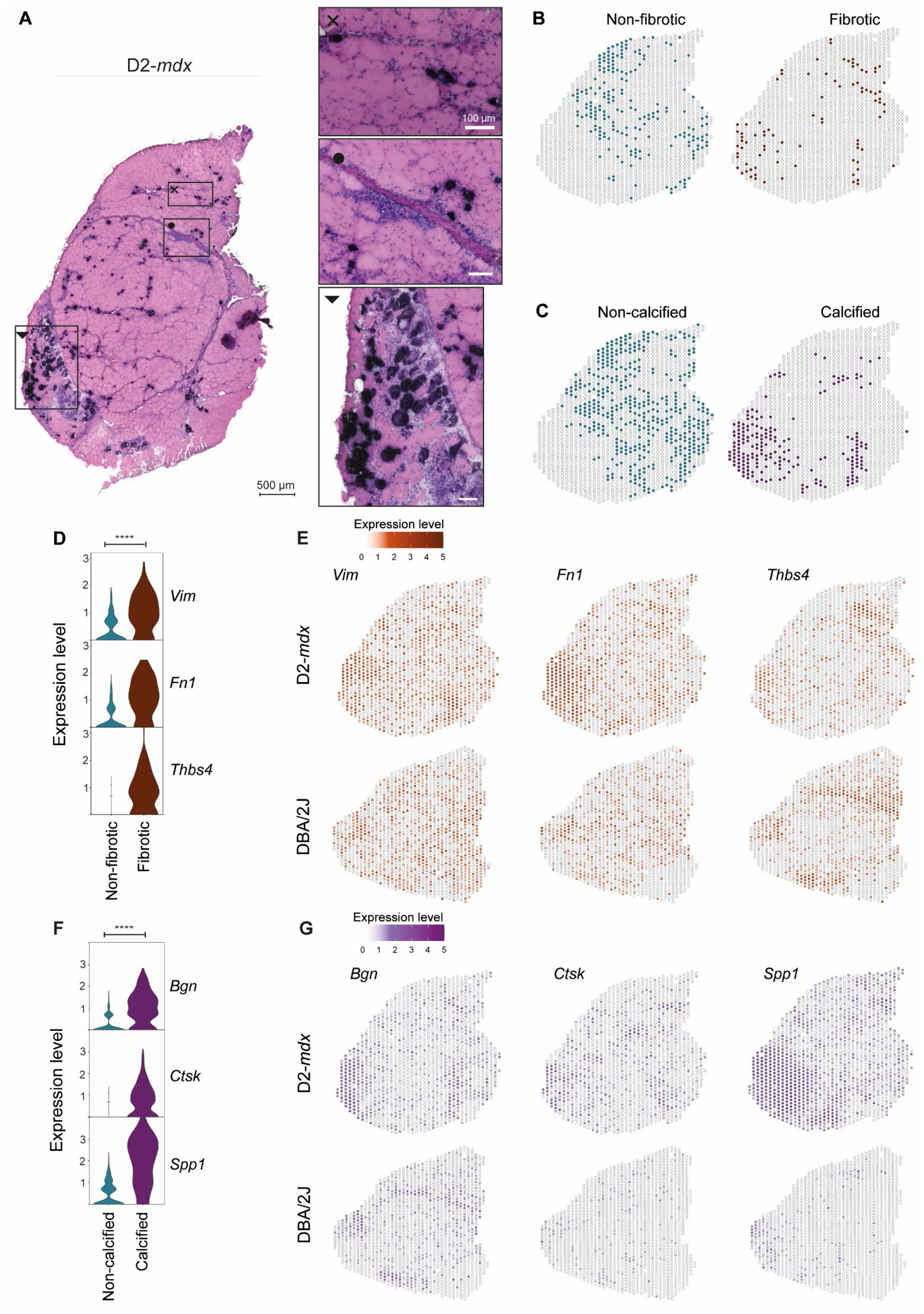
Differential expression analysis in *D2-mdx* calcified and fibrotic tissue displays upregulated molecular markers. (A) HE stained D2-mdx QUA sample with three zoomed-in areas, × displaying fibrotic infiltration between the myofibers, • showing part of the connective tissue sheet with surrounding inflammation, calcification and fibrotic infiltration and ▾ displaying the most severely affected area of the muscle section with extensive calcification, inflammation and necrosis (B) Selected spots belonging to the categories “non-fibrotic” or “fibrotic” as well as (C) “non-calcified” and “calcified” were included in the differential gene expression analysis (D) Violin plot showing expression levels of selected genes Vim, Fn1 and Thbs4 which were significantly upregulated in the “fibrotic” spots (E) Spatially plotted expression of Vim, Fn1 and Thbs4 in D2-mdx and DBA/2J muscle (**** representing a p-value <0.0001) (F) Expression levels of selected genes Bgn, Ctsk and Spp1 which were significantly upregulated in the “calcified” spots compared to the “non-calcified” spots (**** representing a p-value <0.0001) (G) Expression of the selected genes Bgn, Ctsk and Spp1 spatially plotted in the D2-mdx and its genetic background matching DBA/2J wildtype.

Comparison of calcified and non-calcified spots was performed to identify genes associated with calcification. Spots belonging to the cluster ‘inflamed and/or calcified fibers’ and expressing calcification marker *Mgp* were considered calcified, while spots in the muscle fibers cluster lacking *Mgp* expression were considered non-calcified (Figure 4C). Among the top 50 calcification markers (summarized in Table S7), we identified *Bgn, Ctsk* and *Spp1*. Upregulation of *Bgn* was seen in areas that are highly affected in the D2-*mdx* muscle (log_2_(FC) = 1.39, adjusted p-value =1.44e-38, Figure 4F-G). *Bgn*, encodes for the ECM proteoglycan biglycan and is therefore also expressed in the connective tissue of healthy muscle sample (Figure 4G). *Bgn* expression mapped to connective tissue in DBA/2J muscle butbut not in the D2-*mdx* sample (Figure 4G), hinting towards changes to the ECM. *Ctsk* was more specifically expressed in calcified areas (log_2_(FC) = 1.23, adjusted p-value = 1.84e-29, Figure 4F-G) with little-to-no expression in the healthy DBA/2J sample (Figure 4G-DBA/2J). *Ctsk* encodes for cathepsin K and is used as a marker for osteoclasts and known to be involved in osteogenesis ^48,49^. Finally, *Spp1* was highly upregulated in the most severely affected region of the D2-*mdx* muscle section (log_2_(FC) = 2.66, adjusted p-value = 5.78e-50, Figure 4F-G). *Spp1* is a marker for osteoblasts and osteoclasts and can be secreted as a cytokine in inflammatory processes ^50^. It is therefore not only considered as marker of tissue calcification which explains the high expression throughout the D2-*mdx* affected areas.

### RNA velocity reveals patterns of differentiation in severely affected skeletal muscle

To assess whether the histopathological changes observed in dystrophic muscle are associated with cellular transcriptional reprogramming, we applied RNA velocity on the *D2-mdx* spatial datasets, which displayed the highest histological variation compared to the wildtype counterpart (Figure 5A) counterpart.

**Figure 5.**
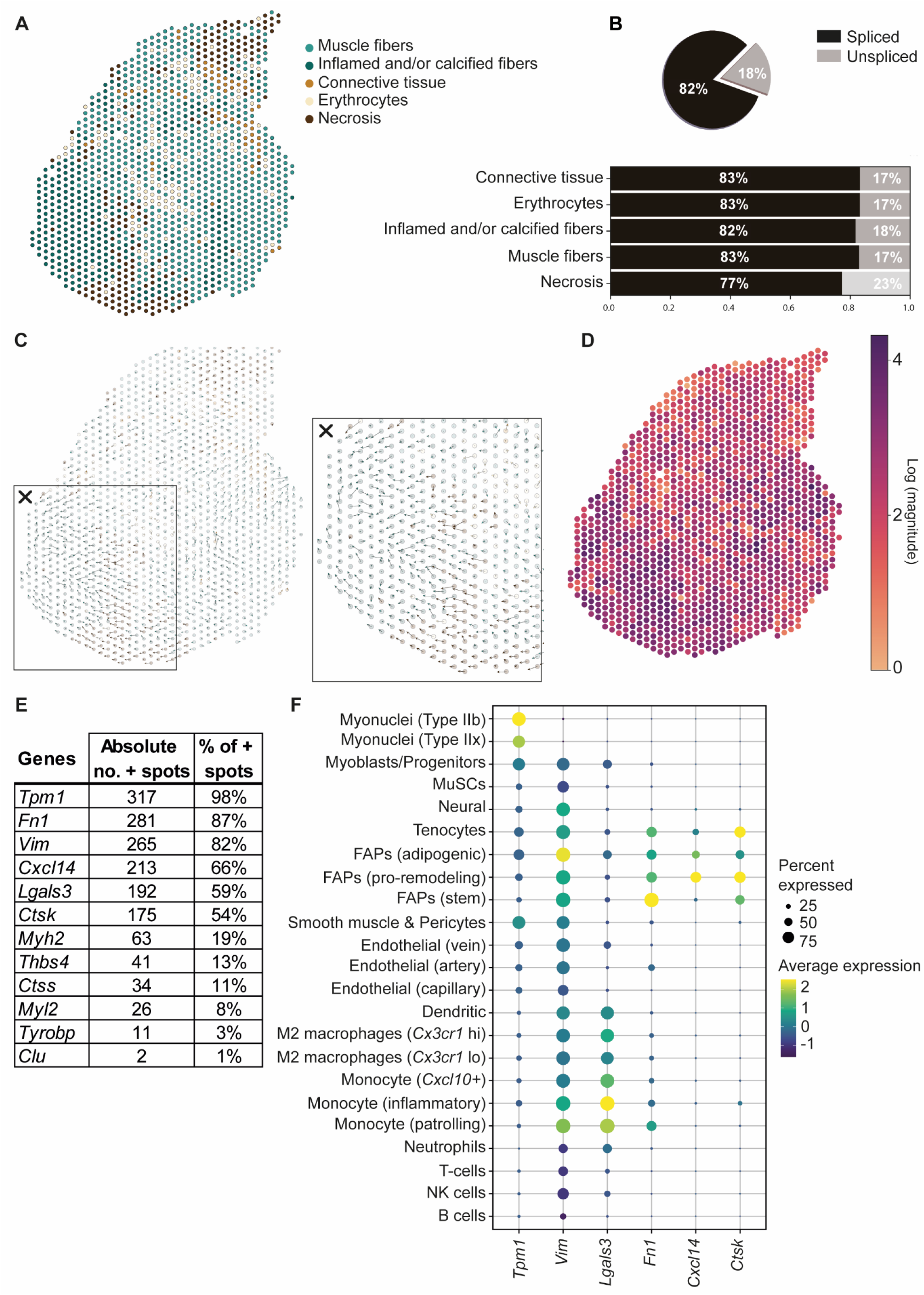
RNA velocity applied on *D2-mdx* muscle shows differentiation patterns in severely affected muscle tissue which is driven by known cell types such as FAPs and macrophages. (A) Annotated clusters of D2-mdx as described before (B) Proportion of spliced/unspliced counts in the D2-mdx sample per annotated cluster (C) Spatial spot-level RNA velocity vectors showing clear differentiation at the severely affected spatial region (× box) (D) Magnitude of the RNA velocity at each spot (E) Summary of the top contributing genes (absolute count and percentage of positive spots) to the differentiation patterns of the severely affected spots (Inflamed and/or calcified fibers). (F) Expression of these top genes in cell types from a reference dataset^52^.

RNA velocity is defined as the time derivative of gene expression in single cells and has been developed to resolve the potential future state of a cell. RNA velocity recovers information about the direction of a single cell in the transcriptional space by distinguishing between newly transcribed pre-mRNAs (unspliced) and mature mRNAs (spliced), which can be detected in sequencing data ^31^. For this analysis we considered each spot on the Visium slide as a single cell. The proportions of reads mapping to spliced and unspliced transcripts (17-23%) fell into the expected range of bulk and single cell RNA sequencing data, with a remarkably higher percentage (23%) of reads mapping to unspliced transcripts in the “necrosis” cluster compared to the other clusters (Figure 5B) ^51^. The proportion of unspliced transcripts was lower in the DBA/2J mouse compared to the *D2-mdx* mouse (Figure S6A-B). The resulting velocities are displayed on the tissue slide as arrows where indicating the direction and strength of change in transcriptional the state of each spot (Figure 5C). We found the strongest pattern in spots surrounding the area with inflamed and/or calcified fibers with arrows pointing towards the calcified area (Figure 5C×-D). This indicates that cells underlying these spots are transitioning likely developing a transcriptionally similar profile to those inflamed and/or calcified fibers. These patterns were not present in the healthy DBA/2J muscle section are very short and (Figure S6C).

We then sought to identify the genes contributing the most to the estimated RNA velocities. Based on the velocity estimated from the spots overlapping with inflamed and/or calcified fibers (324 spots), we identified genes within the top 5 contributing genes in each spot (see Methods for details) (Figure 5E). Next, we checked the cell-type specific expression of those highly contributing genes using the reference dataset of McKellar and colleagues (Figure 5F) ^52^. While the analysis showed that myonuclei and other cell types can participate to the directional pattern by transcribing genes such as *Tpm1* and *Vim*, macrophages and monocytes participated by expressing *Lgals3*, and FAPs contributed with the expression of *Fn1*, *Cxcl14* and *Ctsk*.

## Discussion

Dystrophic muscles are exposed to chronic muscle damage, which eventually leads to replacement of muscle fibers by fibrotic and adipose tissues ^11^. The presence of cell-types and gene expression signatures have been shown to play key-roles in DMD pathogenesis. So far, it has thus far been impossible to directly link the histological alterations to transcriptional changes. Here we used spatial transcriptomics to overcome this limitation. We identify genes and cell types associated with dystrophic features such as muscle regeneration, fibrosis and calcification, clarifying how FAPs, macrophages and myonuclei participate in the transition of muscle mass into fibro-adipogenic tissue.

Based on the direct link between histology and gene expression in the Visium data, we validated some known biomarkers of muscle regeneration as well as revealed previously unknown associations. *Myl4* is a known marker for regeneration and it has been previously suggested to be a useful marker for monitoring dystrophic changes ^53^. *Sparc* is a less known regeneration marker, but upregulation of the protein product SPARC has been shown to reflect the severity of lesions in DMD and Becker muscular dystrophy biopsies ^41,54^. Extensive upregulation of *Sparc* may have an adverse effect on muscle regeneration capabilities ^40^. Additionally, Upregulation of extracellular matrix genes (including *Sparc*) in DMD muscle has also been associated with the dystrophic changes in the muscle such as necrosis, inflammation and muscle regeneration ^53^. Although *Sparc* was identified as a regeneration marker in the *mdx* mouse, we observed high levels of *Sparc* expression in the wildtype (C57BL10) sample. *Sparc* is thought to have contradicting roles depending on its localization. Intracellularly, *Sparc* has a mechanistic role modulating the cytoskeletal structure and function in skeletal muscle ^55^. Extracellularly, the calcium-binding glycoprotein SPARC, also known as osteonectin, encoded by the *Sparc* gene regulates cell interaction with the extracellular matrix ^53,56^. Because of this extracellular activity, high expression levels of *Sparc* expression in healthy skeletal muscle, especially in the connective tissue sheet, are less surprising. Finally, we identified *Hspg2* as a biomarker for muscle regeneration. The *Hspg2* gene encodes perlecan, a heparan sulfate proteoglycan suggested to encompass several functions in cell growth, tissue organization and differentiation ^57^. Perlecan has not been linked to and muscular dystrophy before, but our data suggest that it could potentially be used to study regeneration in dystrophic muscle. What is known however, is that a deficiency of perlecan attenuates skeletal muscle atrophy ^58^. Indeed, upregulation of *Hspg2* was observed in the necrosis cluster where atrophic muscle fibers were present. Further studies are needed to understand the role of *Hspg2* in DMD pathology.

In the severely affected D2-*mdx* mouse, pathological changes to the dystrophic muscle show partial co-localization, which makes it challenging to identify biomarkers specifically related to one of these histopathological changes. Spatial transcriptomics allowed us to select areas/spots where specific pathological changes are exclusively present, enabling the identification of genes specific for either fibrosis or calcification. *Fn1* and *Vim* are well-established fibroblast markers and this study underlines this by re-identifying them as important biomarkers of fibrosis in the D2-*mdx* mouse model. *Fn1* has been used as a target for therapeutic interventions aiming to reduce fibrotic infiltration in DMD pathology ^26,59^. The selected fibrotic markers seem to play important roles in ECM construction and remodeling, which is in line with previous literature where staining of *Vim* has been used and described as a marker for profound ECM changes in dystrophic muscles ^60^. Expression of *Thbs4* was predominantly linked to the perimysium in wildtype mice. *Thbs4* encodes thrombospondin-4 (TSP-4), which is thought to function as a structural protein in tendons and connective tissue ^61–63^. However, *Thbs4* was found to be upregulated in fibrotic regions. In a previous study, TSP-4 overexpression in a dystrophic Drosophila model, rescued the phenotype ^64^. This may be due to the fact that TSP-4 helps in stabilizing the sarcolemma in skeletal myofibers and regulates the production and composition of the ECM ^64^. The upregulation of *Thbs4* in the assigned fibrotic spots in our D2-*mdx* could either be explained by a compensatory mechanism aiming to stabilize muscle fibers, or to a fibrotic / ECM remodeling phenotype.

*Bgn* is a marker for fibrosis previously described in *mdx* mice and DMD patients ^65,66^. More specifically, injection of recombinant human biglycan (rhBGN) has been proposed as a potential therapeutic intervention for DMD following promising studies in *mdx* mice ^67^. In other diseases, upregulation of *Bgn* has been linked to contribute to a pro-osteogenic effect in vascular calcification ^68,69^. Moreover, *Bgn*-deficient mice show reduced growth, decreased bone mass and osteoporosis ^70^. We think that the role of *Bgn* might not be limited to be the fibrotic process but rather to contribute to the calcification/ossification observed in *D2-mdx* mice. On the other hand, the spatial distribution of *Ctsk* expression, which has been used as efficacy outcome after therapeutic intervention aiming to decrease dystrophic calcification ^49^, has shown to be very specifically expressed in these calcified myofibers in the *D2-mdx* sample, suggesting that *Ctsk* could be a more specific marker of calcification/ossification than *Bgn*. Finally, osteopontin, the product of the *Spp1* gene, has been linked to DMD pathology in different ways. It has been shown that a SNP at the promoter site is associated with disease severity and it has been shown to affect regeneration and inflammation levels ^50,71,72^. Another study linked *Spp1* upregulation to vascular smooth muscle cell calcification, supporting the association of this gene in the calcified areas of the *D2-mdx* mouse ^73^.

While the current limitation in the spatial resolution of the Visium platform to the 100μm spot size, we could estimate the cell type contribution using a deconvolution strategy based on existing snRNAseq data. This reference dataset, however, was not from the same muscle type or mouse model, and therefore did not perfectly match our spatial data. The deconvolution results indicated that multiple cell types (mainly myonuclei) are contributing to the expression profile of the Visium spots and further support the annotations we assigned to the spot clusters.

Our proof-of-principle study (including *n* = 1 per mouse model) allowed us to find and link gene expression profiles to clusters that reflected the histology of our skeletal muscle samples. This study shows that the spatial transcriptomics approach is feasible for muscular dystrophies. It shows how it adds spatial information compared to single-cell approaches, and high-throughput compared to targeted gene specific approaches. The spatial approach we took shows great promise for the identification of biomarkers behind histopathological changes observed in dystrophic muscle. Identifying such biomarkers has the potential to enable the identification of therapeutic targets and evaluate the effects of treatments on histological alterations.

## Authorship contribution statement

L.G.M. Heezen carried out the experiments, analysis and drafted the first version of the manuscript. T. Abdelaal contributed by applying RNA velocity on the spatial datasets and writing the methods on this analysis. P. Spitali, A. Mahfouz, M. van Putten and A. Aartsma-Rus have jointly supervised the work. All authors provided feedback and comments.

## References

1. Mendell, J. R. & Lloyd-Puryear, M. Report Of MDA Muscle Disease Symposium On Newborn Screening For Duchenne Muscular Dystrophy. Muscle and Nerve 48, 21–26 (2013).

2. Bladen, C. L. et al. The TREAT-NMD DMD global database: Analysis of more than 7,000 duchenne muscular dystrophy mutations. Hum. Mutat. 36, 395–402 (2015).

3. Hoffman, E. P., Brown, R. H. & Kunkel, L. M. Dystrophin: The protein product of the duchenne muscular dystrophy locus. Cell 51, 919–928 (1987).

4. Le Rumeur, E., Winder, S. J. & Hubert, J. F. Dystrophin: More than just the sum of its parts. Biochim. Biophys. Acta - Proteins Proteomics 1804, 1713–1722 (2010).

5. Ervasti, J. M. Dystrophin, its interactions with other proteins, and implications for muscular dystrophy. Biochim. Biophys. Acta - Mol. Basis Dis. 1772, 108–117 (2007).

6. Gao, Q. & McNally, E. The Dystrophin Complex: structure, function and implications for therapy. Compr Physiol 5, 1223–1239 (2015).

7. Petrof, B. J., Shrager, J. B., Stedman, H. H., Kelly, A. M. & Sweeney, H. L. Dystrophin protects the sarcolemma from stresses developed during muscle contraction. Proc. Natl. Acad. Sci. U. S. A. 90, 3710–3714 (1993).

8. Ogura, Y., Tajrishi, M. M., Sato, S., Hindi, S. M. & Kumar, A. Therapeutic potential of matrix metalloproteinases in Duchenne muscular dystrophy. Front. Cell Dev. Biol. 2, 1–11 (2014).

9. Cirak, S. et al. Exon skipping and dystrophin restoration in patients with Duchenne muscular dystrophy after systemic phosphorodiamidate morpholino oligomer treatment: An open-label, phase 2, dose-escalation study. Lancet 378, 595–605 (2011).

10. Zacharias, J. M. & Anderson, J. E. Muscle regeneration after imposed injury is better in younger than older mdx dystrophic mice. J. Neurol. Sci. 104, 190–196 (1991).

11. Klingler, W., Jurkat-Rott, K., Lehmann-Horn, F. & Schleip, R. The role of fibrosis in Duchenne muscular dystrophy. Acta Myol. 31, 184–195 (2012).

12. Emery, A. E. The muscular dystrophies. Lancet 359, 687–695 (2002).

13. Guiraud, S. & Davies, K. E. Pharmacological advances for treatment in Duchenne muscular dystrophy. Curr. Opin. Pharmacol. 34, 36–48 (2017).

14. Kharraz, Y., Guerra, J., Pessina, P., Serrano, A. L. & Muñoz-Cánoves, P. Understanding the process of fibrosis in duchenne muscular dystrophy. Biomed Res. Int. 2014, (2014).

15. Forcina, L., Pelosi, L., Miano, C. & Musarò, A. Insights into the pathogenic secondary symptoms caused by the primary loss of dystrophin. J. Funct. Morphol. Kinesiol. 2, (2017).

16. Capitanio, D. et al. Comparative proteomic analyses of Duchenne muscular dystrophy and Becker muscular dystrophy muscles: changes contributing to preserve muscle function in Becker muscular dystrophy patients. J. Cachexia. Sarcopenia Muscle 11, 547–563 (2020).

17. Pescatori, M. et al. Gene expression profiling in the early phases of DMD: a constant molecular signature characterizes DMD muscle from early postnatal life throughout disease progression. FASEB J. 21, 1210–1226 (2007).

18. Coley, W. D. et al. Effect of genetic background on the dystrophic phenotype in mdx mice. Hum. Mol. Genet. 25, 130–145 (2016).

19. van Putten, M. et al. Natural disease history of the D2-mdx mouse model for Duchenne muscular dystrophy. FASEB J. 33, 8110–8124 (2019).

20. Gordish-Dressman, H. et al. ‘of mice and measures’: A project to improve how we advance duchenne muscular dystrophy therapies to the clinic. J. Neuromuscul. Dis. 5, 407–417 (2018).

21. Turk, R. et al. Muscle regeneration in dystrophin-deficient mdx mice studied by gene expression profiling. BMC Genomics 6, 1–15 (2005).

22. Marotta, M. et al. Muscle genome-wide expression profiling during disease evolution in mdx mice. Physiol. Genomics 37, 119–32 (2009).

23. Brinkmeyer-Langford, C. et al. Expression profiling of disease progression in canine model of Duchenne muscular dystrophy. PLoS One 13, 1–21 (2018).

24. van Putten, M. et al. Comparison of skeletal muscle pathology and motor function of dystrophin and utrophin deficient mouse strains. Neuromuscul. Disord. 22, 406–417 (2012).

25. Delaney, K., Kasprzycka, P., Ciemerych, M. A. & Zimowska, M. The role of TGF-β1 during skeletal muscle regeneration. Cell Biol. Int. 41, 706–715 (2017).

26. Zanotti, S., Gibertini, S., Savadori, P., Mantegazza, R. & Mora, M. Duchenne muscular dystrophy fibroblast nodules: A cell-based assay for screening anti-fibrotic agents. Cell Tissue Res. 352, 659–670 (2013).

27. Ismaeel, A. et al. Role of transforming growth factor-β in skeletal muscle fibrosis: A review. International Journal of Molecular Sciences vol. 20 (2019).

28. Morales, M. G., Acuña, M. J., Cabrera, D., Goldschmeding, R. & Brandan, E. The pro-fibrotic connective tissue growth factor (CTGF/CCN2) correlates with the number of necrotic-regenerative foci in dystrophic muscle. J. Cell Commun. Signal. 12, 413–421 (2018).

29. Natarajan, A., Lemos, D. R. & Rossi, F. M. V. Fibro/adipogenic progenitors: A double-edged sword in skeletal muscle regeneration. Cell Cycle 9, 2045–2046 (2010).

30. Moratal, C., Arrighi, N., Dechesne, C. A. & Dani, C. Control of muscle fibro-adipogenic progenitors by myogenic lineage is altered in aging and Duchenne muscular dystrophy. Cell. Physiol. Biochem. 53, 1029–1045 (2019).

31. La Manno, G. et al. RNA velocity of single cells. Nature 560, 494–498 (2018).

32. McInnes, L., Healy, J. & Melville, J. UMAP: Uniform Manifold Approximation and Projection for Dimension Reduction. (2018).

33. Elosua-Bayes, M., Nieto, P., Mereu, E., Gut, I. & Heyn, H. SPOTlight: Seeded NMF regression to deconvolute spatial transcriptomics spots with single-cell transcriptomes. Nucleic Acids Res. 49, E50 (2021).

34. Rodriques, S. G. et al. Slide-seq: A scalable technology for measuring genome-wide expression at high spatial resolution. Science (80-.). 363, 1463–1467 (2019).

35. Chemello, F. et al. Degenerative and regenerative pathways underlying Duchenne muscular dystrophy revealed by single-nucleus RNA sequencing. Proc. Natl. Acad. Sci. U. S. A. 117, 29691–29701 (2020).

36. Melsted, P. et al. Modular, efficient and constant-memory single-cell RNA-seq preprocessing. Nat. Biotechnol. 39, 813–818 (2021).

37. Bray, N. L., Pimentel, H., Melsted, P. & Pachter, L. Near-optimal probabilistic RNA-seq quantification. Nat. Biotechnol. 34, 525–527 (2016).

38. Bergen, V., Lange, M., Peidli, S., Wolf, F. A. & Theis, F. J. Generalizing RNA velocity to transient cell states through dynamical modeling. Nat. Biotechnol. 38, 1408–1414 (2020).

39. Coulton, G. R., Morgan, J. E., T. A. Partridge & Sloper, J. C. THE mdx MOUSE SKELETAL MUSCLE MYOPATHY: I. A HISTOLOGICAL, MORPHOMETRIC AND BIOCHEMICAL INVESTIGATION. Neuropathol. Appl. Neurobiol. 14, 53–70 (1988).

40. Petersson, S. J. et al. SPARC is up-regulated during skeletal muscle regeneration and inhibits myoblast differentiation. Histol. Histopathol. 28, 1451–1460 (2013).

41. Jørgensen, L. H. et al. Secreted protein acidic and rich in cysteine (SPARC) in human skeletal muscle. J. Histochem. Cytochem. 57, 29–39 (2009).

42. Sato, K. et al. Improvement of muscle healing through enhancement of muscle regeneration and prevention of fibrosis. Muscle and Nerve 28, 365–372 (2003).

43. Sosa, P. et al. Aging-related hyperphosphatemia impairs myogenic differentiation and enhances fibrosis in skeletal muscle. J. Cachexia. Sarcopenia Muscle 12, 1266–1279 (2021).

44. Sun, Y. et al. miR-24 and miR-122 Negatively Regulate the Transforming Growth Factor-β/Smad Signaling Pathway in Skeletal Muscle Fibrosis. Mol. Ther. - Nucleic Acids 11, 528–537 (2018).

45. Zhang, R., Lv, L., Ban, W., Dang, X. & Zhang, C. Identification of Hub Genes in Duchenne Muscular Dystrophy: Evidence from Bioinformatic Analysis. J. Comput. Biol. 27, 1–8 (2020).

46. Kallenbach, J. G. et al. Muscle-specific functional deficits and lifelong fibrosis in response to paediatric radiotherapy and tumour elimination. J. Cachexia. Sarcopenia Muscle (2022) doi:10.1002/jcsm.12902.

47. Contreras, O. et al. Cross-talk between TGF-β and PDGFRα signaling pathways regulates the fate of stromal fibro-adipogenic progenitors. J. Cell Sci. 132, (2019).

48. Troen BR. The role of cathepsin K in normal bone resorption. Drug News Perspect. 17, 19–28 (2004).

49. Bauer, C. et al. Etidronate prevents dystrophic cardiac calcification by inhibiting macrophage aggregation. Sci. Rep. 8, 1–13 (2018).

50. Bello, L. et al. Importance of SPP1 genotype as a covariate in clinical trials in Duchenne muscular dystrophy. Neurology 79, 159–162 (2012).

51. Picelli, S. et al. Smart-seq2 for sensitive full-length transcriptome profiling in single cells. Nat. Methods 10, 1096–1100 (2013).

52. McKellar, D. W. et al. Large-scale integration of single-cell transcriptomic data captures transitional progenitor states in mouse skeletal muscle regeneration. Commun. Biol. 4, 1–12 (2021).

53. Noguchi, S. et al. cDNA microarray analysis of individual Duchenne muscular dystrophy patients. Hum. Mol. Genet. 12, 595–600 (2003).

54. Haslett, J. N. et al. Gene expression comparison of biopsies from Duchenne muscular dystrophy (DMD) and normal skeletal muscle. Proc. Natl. Acad. Sci. U. S. A. 99, 15000–15005 (2002).

55. Jørgensen, L. H. et al. SPARC Interacts with Actin in Skeletal Muscle in Vitro and in Vivo. Am. J. Pathol. 187, 457–474 (2017).

56. Brekken, R. A. & Sage, E. H. SPARC, a matricellular protein: at the crossroads of cell-matrix communication. Matrix Biol. 19, 816–827 (2001).

57. Hassel, J., Yoshihiko, Y. & Arikawa-Hirasawa, E. Role of perlecan in development and diseases. Glycoconj. J. 19, 263–267 (2003).

58. Nakada, S., Yamashita, Y., Machida, S., Miyagoe-Suzuki, Y. & Arikawa-Hirasawa, E. Perlecan Facilitates Neuronal Nitric Oxide Synthase Delocalization in Denervation-Induced Muscle Atrophy. Cells 9, (2020).

59. Piñol-Jurado, P. et al. Nintedanib decreases muscle fibrosis and improves muscle function in a murine model of dystrophinopathy. Cell Death Dis. 9, (2018).

60. Holland, A., Murphy, S., Dowling, P. & Ohlendieck, K. Pathoproteomic profiling of the skeletal muscle matrisome in dystrophinopathy associated myofibrosis. Proteomics 16, 345–366 (2016).

61. Södersten, F., Ekman, S., Schmitz, M., Paulsson, M. & Zaucke, F. Thrombospondin-4 and cartilage oligomeric matrix protein form heterooligomers in equine tendon. Connect. Tissue Res. 47, 85–91 (2006).

62. Hauser, N., Paulsson, M., Kale, A. A. & DiCesare, P. E. Tendon extracellular matrix contains pentameric thrombospondin-4 (TSP-4). FEBS Lett. 368, 307–310 (1995).

63. Jelinsky, S. A., Archambault, J., Li, L. & Seeherman, H. Tendon-selective genes identified from rat and human musculoskeletal tissues. J. Orthop. Res. 28, 289–297 (2010).

64. Vanhoutte, D. et al. Thrombospondin expression in myofibers stabilizes muscle membranes. Elife 5, 1–33 (2016).

65. Zanotti, S. et al. Decorin and biglycan expression is differentially altered in several muscular dystrophies. Brain 128, 2546–2555 (2005).

66. ‘t Hoen, P. A. C. et al. Gene expression profiling to monitor therapeutic and adverse effects of antisense therapies for Duchenne muscular dystrophy. Pharmacogenomics 7, 281–297 (2006).

67. Amenta, A. R. et al. Biglycan recruits utrophin to the sarcolemma and counters dystrophic pathology in mdx mice. Proc. Natl. Acad. Sci. U. S. A. 108, 762–767 (2011).

68. Tsang, H. G. et al. Expression of Calcification and Extracellular Matrix Genes in the Cardiovascular System of the Healthy Domestic Sheep (Ovis aries). Front. Genet. 11, (2020).

69. Song, R. et al. Biglycan induces the expression of osteogenic factors in human aortic valve interstitial cells via toll-like receptor-2. Arterioscler. Thromb. Vasc. Biol. 32, 2711–2720 (2012).

70. Xu, T. et al. Targeted disruption of the biglycan gene leads to an osteoporosis-like phenotype in mice. Nat. Genet. 20, 78–82 (1998).

71. Vianello, S. et al. SPP1 genotype and glucocorticoid treatment modify osteopontin expression in Duchenne muscular dystrophy cells. Hum. Mol. Genet. 26, 3342–3351 (2017).

72. Kyriakides, T. et al. SPP1 genotype is a determinant of disease severity in Duchenne muscular dystrophy: Predicting the severity of duchenne muscular dystrophy: Implications for treatment. Neurology 77, 1858–1859 (2011).

73. Patel, J. J. et al. Differing calcification processes in cultured vascular smooth muscle cells and osteoblasts. Exp. Cell Res. 380, 100–113 (2019).

