## Supplementary figures and tables for "Spatial transcriptomics reveal markers of histopathological changes in Duchenne muscular dystrophy mouse models"

### Supplementary information

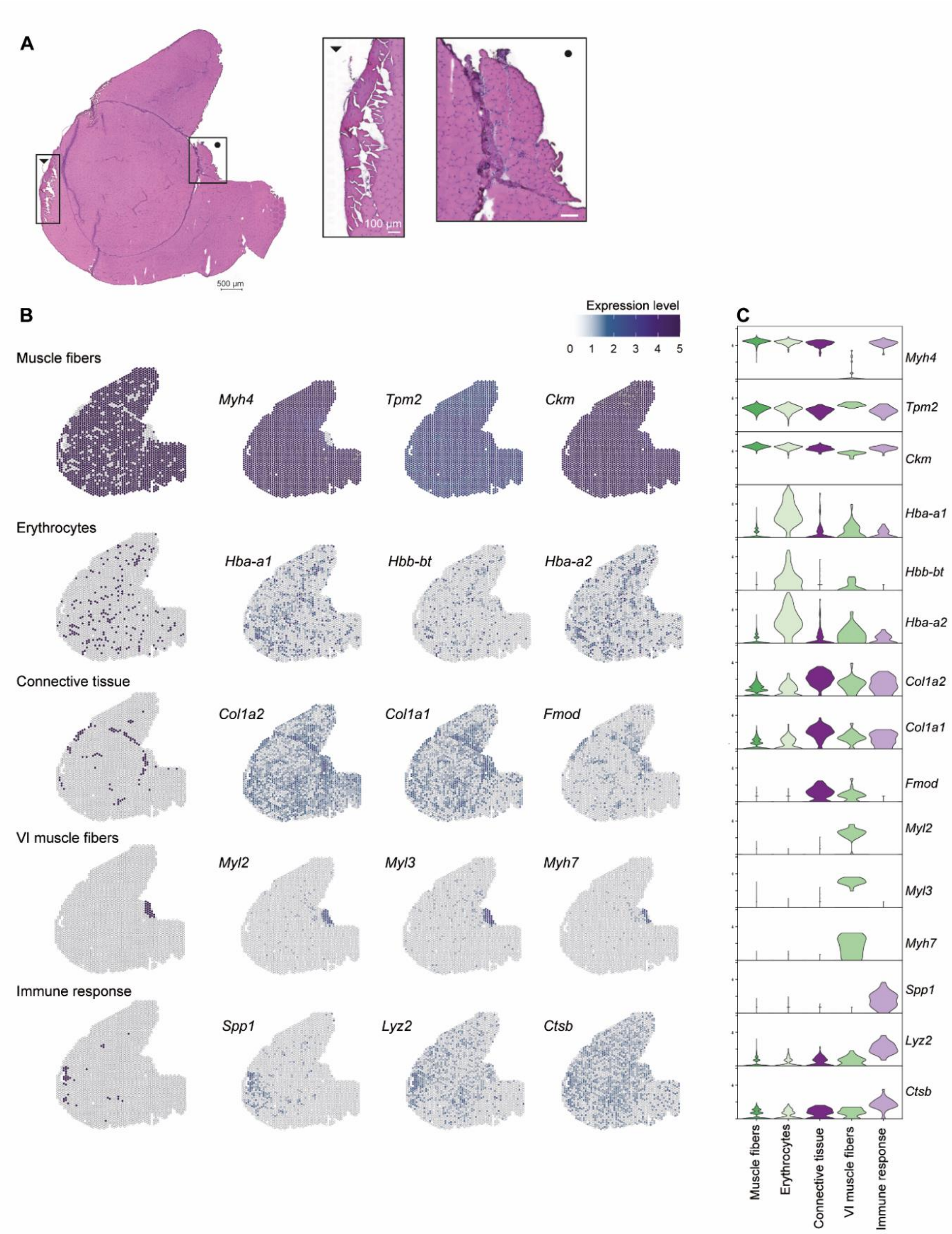


*Figure S1.* **Annotated clusters of the C57BL10 muscle based on histological features and the upregulated expression of marker genes in these clusters.** *(A) HE stained C57BL10 QUA sample with two zoomed-in areas, ▼ displaying damaged tissue area close to the immune response cluster, • showing VI muscle (B) All annotated clusters spatially plotted with the main marker genes and their gene expression level throughout the tissue section (C) Violin plot showing the expression level of the marker genes across the annotated clusters.*

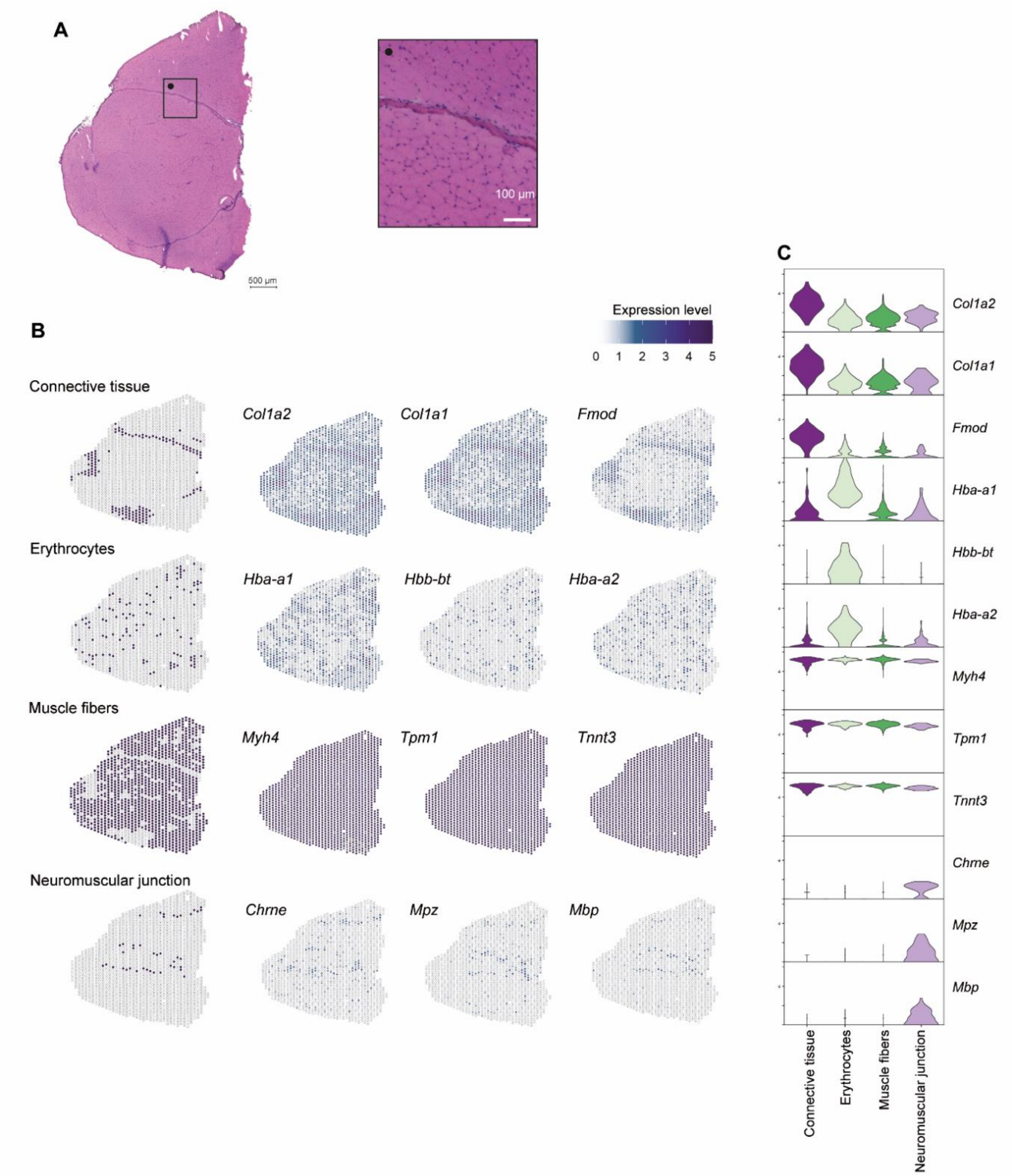


*Figure S2.* **Annotated clusters of the DBA/2J muscle based on histological features and the upregulated expression of marker genes in these clusters.** *(A) HE stained DBA/2J QUA sample with one zoomed-in area, • showing part of the connective tissue sheet and representative healthy muscle fibers (B) All annotated clusters spatially plotted with the main marker genes and their gene expression level throughout the tissue section (C) Violin plot showing the expression level of the marker genes across the annotated clusters.*

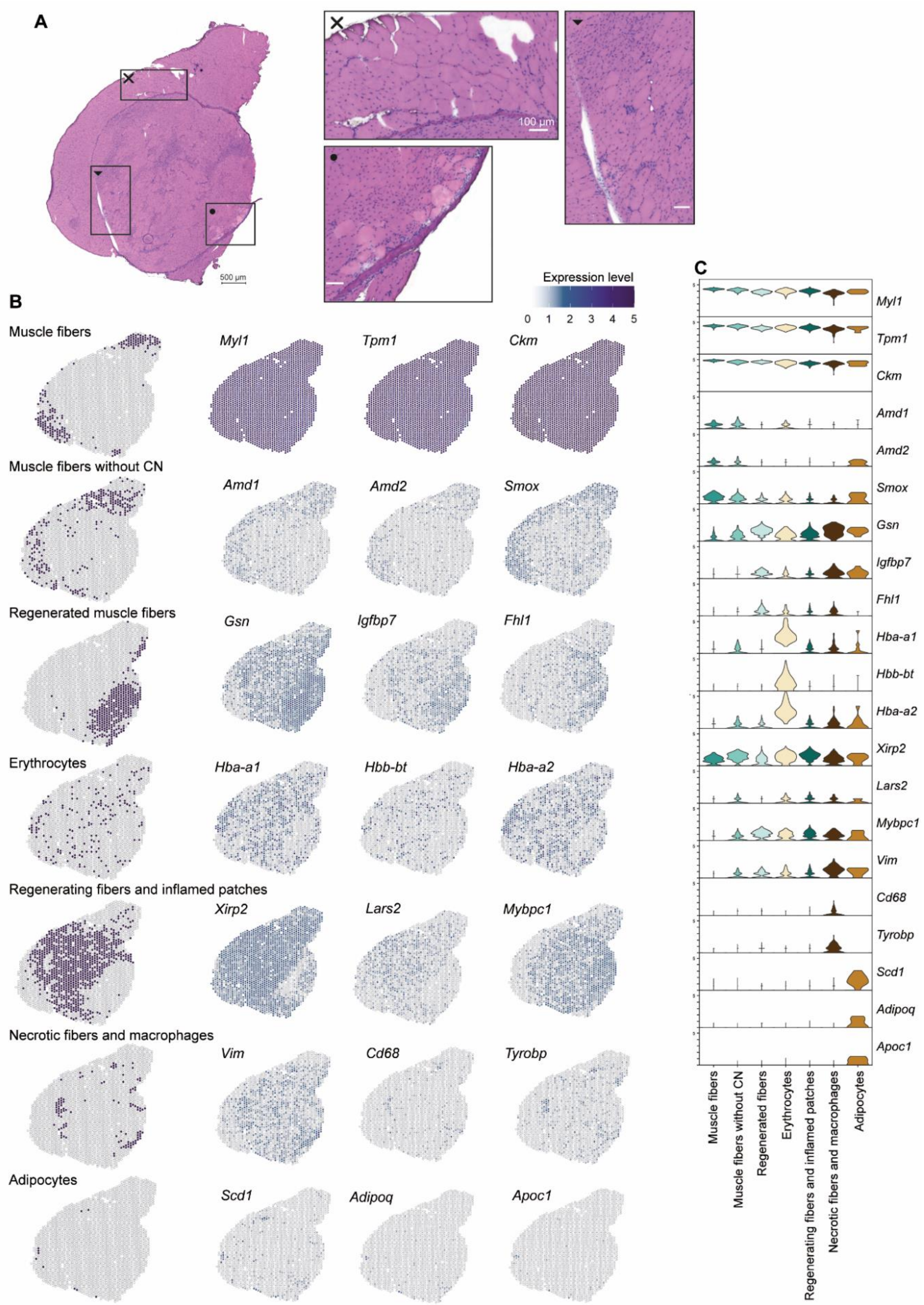


*Figure S3.* **Annotated clusters of the *mdx* muscle based on histological features and the upregulated expression of marker genes in these clusters.** *(A) HE stained mdx QUA sample with three zoomed-in areas, ✕ displaying mature myofibers without central nuclei, • showing some necrotic fibers and lastly ▼ displaying an area of regeneration and mild inflammation (B) All annotated clusters spatially plotted with the main marker genes and their gene expression level throughout the tissue section (C) Violin plot showing the expression level of the marker genes across the annotated clusters.*

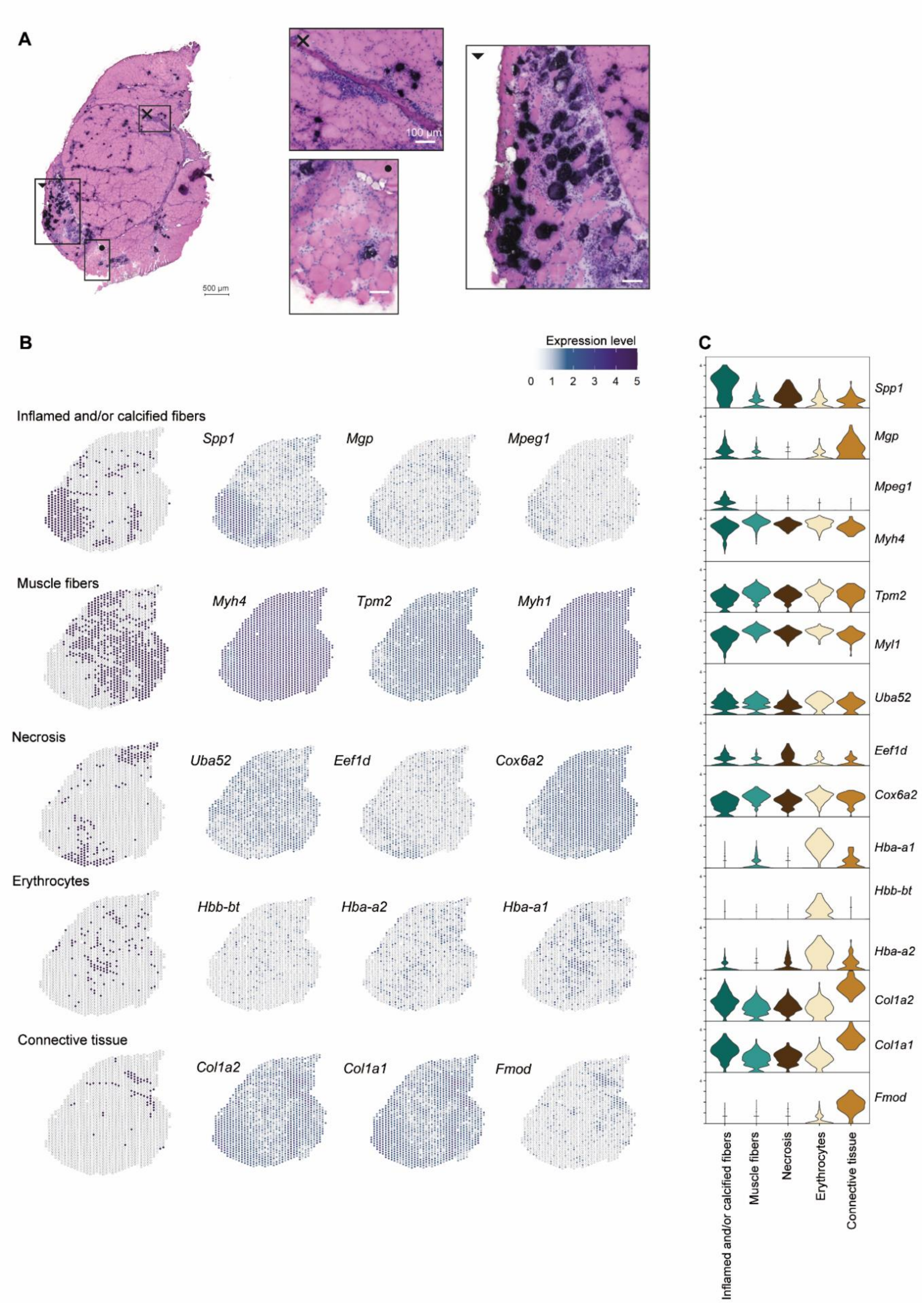


*Figure S4.* **D2-*mdx* annotated clusters based on histological features and the upregulated expression of marker genes in these clusters.** *(A) HE stained D2-mdx QUA sample with three zoomed-in areas, ✕ displaying inflammation around the connective tissue sheet, • showing some necrotic fibers and lastly ▼ displaying the most severely affected area of the muscle section with extensive calcification, inflammation and fibrosis (B) All annotated clusters spatially plotted with the*

*main marker genes and their gene expression level throughout the tissue section (C) Violin plot showing the expression level of the marker genes across the annotated clusters.*

*Table S1. C57BL10 top genes per cluster*

| | Cluster | Gene | p-val | avg log2FC | pct1 | pct2 | p-val adjusted |
|---|---|---|---|---|---|---|---|
| C57BL10 | Muscle fibers | *Myh4* | 1.08e-71 | 0.318 | 1 | 0.990 | 1.40e-67 |
| | | *Ckm* | 1.43e-102 | 1.159 | 0.991 | 0.920 | 1.85e-98 |
| | | *Tnnt3* | 4.69e-123 | 0.355 | 1 | 1 | 6.05e-119 |
| | | *Tpm1* | 4.25e-123 | 0.424 | 1 | 1 | 5.49e119 |
| | Erythrocytes | *Hba-a1* | 1.77e-152 | 4.654 | 0.951 | 0.263 | 2.28e-148 |
| | | *Hbb-bt* | 4.17e-141 | 2.931 | 0.716 | 0.096 | 5.38e-137 |
| | | *Hba-a2* | 1.20e-136 | 4.446 | 0.926 | 0.270 | 1.55e-132 |
| | | *Hbb-bs* | 1.62e-119 | 4.893 | 0.980 | 0.574 | 2.09e-115 |
| | Connective tissue | *Col1a1* | 2.72e-44 | 2.070 | 0.945 | 0.508 | 3.51e-40 |
| | | *Fmod* | 1.19e-54 | 1.483 | 0.769 | 0.177 | 1.53e-50 |
| | | *Col1a2* | 1.48e-41 | 2.000 | 0.967 | 0.649 | 1.91e-37 |
| | | *Thbs4* | 1.20e-43 | 1.269 | 0.703 | 0.172 | 1.55e-39 |
| | Immune response | *Lyz2* | 6.48e-34 | 2.772 | 1 | 0.300 | 8.35e-30 |
| | | *Spp1* | 8.57e-71 | 2.656 | 0.917 | 0.083 | 1.11e-66 |
| | | *Lgals3* | 2.96e-37 | 2.002 | 0.722 | 0.102 | 3.82e-33 |
| | | *Ctsb* | 1.52e-25 | 1.163 | 0.583 | 0.091 | 1.96e-21 |
| | VI muscle fibers | *Myh2* | 6.85e-30 | 4.894 | 1 | 0.227 | 8.84e-26 |
| | | *Myl3* | 4.29e-90 | 4.377 | 1 | 0.053 | 5.53e-86 |
| | | *Myl2* | 3.41e-122 | 3.761 | 0.958 | 0.032 | 4.40e-118 |
| | | *Myh7* | 4.81e-120 | 3.092 | 0.792 | 0.020 | 6.21e-116 |

*Table S2. DBA/2J  top genes per cluster*

| | Cluster | Gene | p-val | avg log2FC | pct1 | pct2 | p-val adjusted |
|---|---|---|---|---|---|---|---|
| DBA/2J | Muscle fibers | *Tpm1* | 1.34e-70 | 0.365 | 1 | 1 | 1.71e-66 |
| | | *Tnnt3* | 1.40e-50 | -0.324 | 1 | 1 | 1.78e-46 |
| | | *Ckm* | 3.28e-62 | 0.273 | 1 | 1 | 4.18e-58 |
| | Neuromuscular junction | *Chrne* | 6.66e-46 | 1.529 | 0.780 | 0.104 | 8.46e-42 |
| | | *Mpz* | 3.09e-38 | 1.792 | 0.634 | 0.074 | 3.93e-34 |
| | | *Pmp22* | 2.70e-16 | 1.472 | 0.659 | 0.189 | 3.43e-12 |
| | | *Mbp* | 6.58e-51 | 1.443 | 0.610 | 0.046 | 8.36e-47 |
| | Erythrocytes | *Hba-a1* | 2.41e-71 | 4.167 | 1 | 0.596 | 3.07e-67 |
| | | *Hbb-bt* | 2.08e-77 | 2.773 | 0.778 | 0.136 | 2.64e-73 |
| | | *Hba-a2* | 5.12e-68 | 2.611 | 0.889 | 0.265 | 6.51e-64 |
| | | *Hbb-bs* | 1.70e-69 | 4.104 | 1 | 0.767 | 2.16e-65 |
| | Connective tissue | *Fmod* | 8.94e-77 | 2.616 | 0.952 | 0.446 | 1.14e-72 |
| | | *Col1a1* | 6.06e-64 | 2.814 | 0.993 | 0.855 | 7.70e-60 |
| | | *Col1a2* | 1.17e-58 | 2.500 | 1 | 0.895 | 1.48e-54 |

| | | Gene | p-val | avg log2FC | pct1 | pct2 | p-val adjusted |
|---|---|---|---|---|---|---|---|
| | | *Sparc* | 1.82e-58 | 1.642 | 1 | 0.945 | 2.32e-54 |

*Table S3. mdx top genes per cluster*

| | Cluster | Gene | p-val | avg log2FC | pct1 | pct2 | p-val adjusted |
|---|---|---|---|---|---|---|---|
| ***mdx*** | **Regenerating fibers with inflamed patches** | *Xirp2* | 1.92e-57 | 0.695 | 0.971 | 0.792 | 2.50e-53 |
| | | *Mybpc1* | 2.04e-22 | 0.601 | 0.831 | 0.523 | 3.08e-39 |
| | | *Lars2* | 9.06e-16 | 0.335 | 0.441 | 0.278 | 1.18e-11 |
| | | *Myh1* | 7.11e-55 | 1.425 | 0.996 | 0.819 | 9.25e-51 |
| | **Muscle fibers** | *Ckm* | 3.03e-90 | 0.553 | 1 | 1 | 3.94e-86 |
| | | *Myl1* | 1.99e-60 | 0.426 | 1 | 1 | 2.60e-56 |
| | | *Tpm1* | 2.01e-43 | 0.331 | 1 | 1 | 2.61e-39 |
| | **Recently regenerated fibers** | *Gsn* | 1.52e-73 | 0.986 | 0.976 | 0.673 | 1.98e-69 |
| | | *Igfbp7* | 1.45e-46 | 0.575 | 0.740 | 0.333 | 1.89e-42 |
| | | *Fhl1* | 8.01e-26 | 0.503 | 0.553 | 0.268 | 1.04e-21 |
| | | *Mybpc1* | 7.91e-25 | 0.478 | 0.825 | 0.558 | 1.03e-20 |
| | **Muscle fibers without CN** | *Amd1* | 6.05e-17 | 0.409 | 0.479 | 0.241 | 7.88e-13 |
| | | *Amd2* | 3.25e-08 | 0.251 | 0.300 | 0.165 | 0.000423 |
| | | *Smox* | 2.80e-09 | 0.290 | 0.612 | 0.425 | 3.64e-05 |
| | | *Nmrk2* | 4.97e-11 | 0.310 | 0.897 | 0.744 | 6.46e-06 |
| | **Erythrocytes** | *Hba-a1* | 1.22e-120 | 4.176 | 1 | 0.371 | 1.58e-116 |
| | | *Hba-a2* | 3.02e-115 | 4.159 | 1 | 0.385 | 3.93e-111 |
| | | *Hbb-bs* | 2.37e-103 | 4.503 | 1 | 0.622 | 5.22e-81 |
| | | *Hbb-bt* | 1.28e-127 | 2.804 | 0.802 | 0.116 | 1.67e-123 |
| | **Necrotic fibers and macrophages** | *Cd68* | 1.04e-24 | 0.582 | 0.299 | 0.052 | 1.35e-20 |
| | | *Vim* | 3.32e-30 | 1.241 | 0.832 | 0.421 | 4.32e-26 |
| | | *Tyrobp* | 1.35e-35 | 0.948 | 0.570 | 0.140 | 1.76e-31 |
| | | *Anxa5* | 9.86e-28 | 0.706 | 0.561 | 0.166 | 1.28e-23 |
| | **Adipocytes** | *Apoc1* | 4.61e-71 | 1.058 | 0.600 | 0.008 | 5.99e-67 |
| | | *Scd1* | 1.00e-26 | 2.697 | 0.900 | 0.069 | 1.30e-22 |
| | | *Cidec* | 2.65e-56 | 1.646 | 0.800 | 0.021 | 3.45e-52 |
| | | *Adipoq* | 5.61e-26 | 1.294 | 0.500 | 0.019 | 7.30e-22 |

*Table S4. D2-mdx top genes per cluster*

| | Cluster | Gene | p-val | avg log2FC | pct1 | pct2 | p-val adjusted |
|---|---|---|---|---|---|---|---|
| **D2-*mdx*** | **Muscle fibers** | *Myh4* | 2.37e-93 | 0.676 | 1 | 1 | 2.92e-89 |
| | | *Tpm1* | 2.55e-65 | 0.515 | 1 | 1 | 3.15e-61 |
| | | *Myl1* | 1.98e-57 | 0.543 | 1 | 0.998 | 2.45e-53 |
| | | *Ckm* | 1.02e-58 | 0.485 | 1 | 1 | 1.26e-54 |
| | **Inflamed and/or calcified fibers** | *Mgp* | 1.86e-13 | 0.4331 | 0.534 | 0.323 | 2.30e-09 |
| | | *Lyz2* | 4.45e-85 | 1.757 | 0.972 | 0.717 | 5.50e-80 |
| | | *Spp1* | 1.89e-85 | 2.460 | 0.904 | 0.557 | 2.34e-81 |
| | | *Mpeg1* | 9.02e-57 | 0.677 | 0.543 | 0.130 | 1.11e-52 |
| | **Necrosis** | *Uba52* | 1.36e-11 | -0.464 | 0.674 | 0.817 | 1.68e-07 |
| | | *Eef1d* | 2.62e-08 | 0.535 | 0.519 | 0.379 | 0.0003237 |
| | | *Cox6a2* | 6.64e-08 | -0.351 | 0.983 | 0.946 | 0.0008203 |
| | | *Vim* | 1.50e-06 | -0.438 | 0.387 | 0.584 | 0.01847 |
| | **Erythrocytes** | *Hba-a1* | 1.10e-88 | 2.770 | 0.977 | 0.311 | 1.36e-84 |
| | | *Hbb-bs* | 2.61e-82 | 3.183 | 1 | 0.489 | 3.22e-78 |
| | | *Hbb-bt* | 8.32e-77 | 1.307 | 0.667 | 0.089 | 1.03e-72 |
| | | *Hba-a2* | 8.40e-55 | 1.846 | 0.829 | 0.272 | 1.04e-50 |
| | **Connective tissue** | *Col1a2* | 2.14e-36 | 2.565 | 1 | 0.901 | 2.64e-32 |
| | | *Col1a1* | 2.14e-35 | 2.361 | 1 | 0.914 | 2.64e-31 |
| | | *Fmod* | 6.65e-54 | 2.334 | 0.951 | 0.230 | 8.21e-50 |
| | | *Chad* | 1.09e-78 | 1.377 | 0.738 | 0.059 | 1.34e-74 |

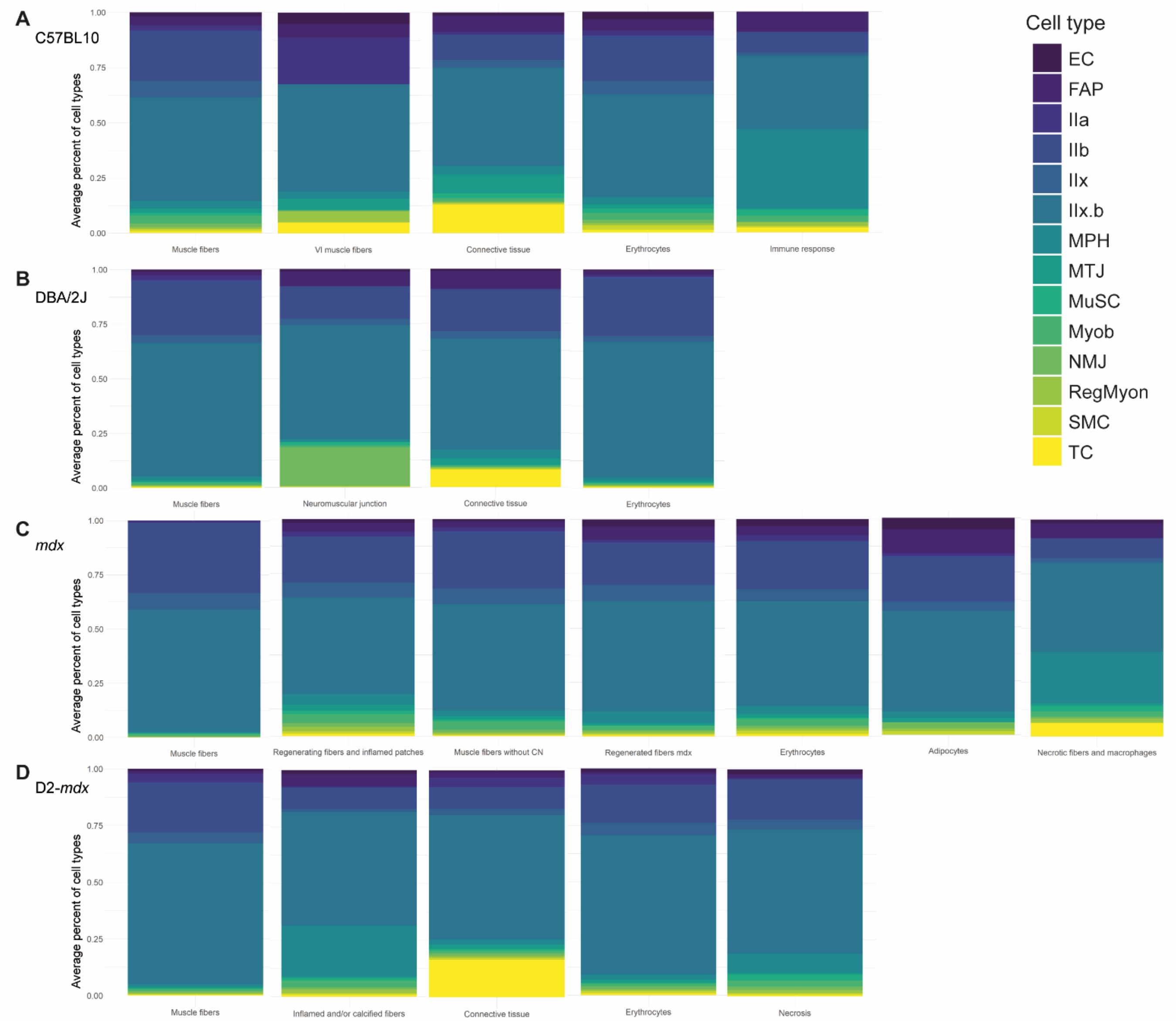


*Figure S5.* **Deconvolution of the spatial data using snRNAseq reference dataset displays average percentage of contributing celltypes to annotated clusters.** *Stacked barplots displaying the average percentage of contributing cell types to the anotated clusters for (A) C57BL10, (B) DBA/2J, (C) mdx and (D) D2-mdx.*

**Differentially expressed genes regeneration.**

*Table S5.* Top 50 regeneration markers ranked on average log2FC.

| Gene | p-value | avg log2FC | pct.1 | pct.2 | Adjusted p-value |
|---|---|---|---|---|---|
| *Myh1* | 7.68e-18 | 1.5136 | 0.962 | 0.776 | 9.99e-14 |
| *Myl4* | 4.23e-21 | 1.4230 | 0.619 | 0.12 | 5.50e-17 |
| *Igfbp7* | 2.60e-54 | 1.3518 | 0.895 | 0 | 3.39e-50 |
| *Col3a1* | 1.39e-25 | 1.3482 | 0.952 | 0.516 | 1.81e-21 |
| *Gsn* | 1.45e-24 | 1.3461 | 0.924 | 0.562 | 1.89e-20 |
| *Fabp4* | 4.43e-21 | 1.2551 | 0.848 | 0.38 | 5.76e-17 |
| *Tmsb10* | 1.70e-16 | 1.2380 | 0.838 | 0.422 | 2.21e-12 |
| *Apoe* | 1.31e-24 | 1.2194 | 0.829 | 0.297 | 1.70e-20 |
| *Casq1* | 1.12e-26 | 1.1213 | 1 | 1 | 1.46e-22 |
| *Actc1* | 2.99e-08 | 1.0753 | 0.952 | 0.964 | 0.00039 |
| *Mgp* | 9.12e-22 | 1.0420 | 0.686 | 0.161 | 1.19e-17 |
| *Mybpc1* | 6.697e-21 | 1.0264 | 0.867 | 0.458 | 8.70e-17 |
| *S100a6* | 2.06e-22 | 1.0197 | 0.924 | 0.427 | 2.68e-18 |
| *Sparc* | 4.31e-20 | 0.8972 | 0.952 | 0.521 | 5.61e-16 |
| *Actn2* | 9.476e-16 | 0.8756 | 0.79 | 0.344 | 1.23e-11 |
| *Lyz2* | 3.10e-17 | 0.8707 | 0.695 | 0.224 | 4.04e-13 |
| *Cd74* | 1.30e-17 | 0.8533 | 0.695 | 0.208 | 1.69e-13 |
| *Hspg2* | 1.35e-19 | 0.8364 | 0.571 | 0.104 | 1.76e-15 |
| *Mb* | 9.27e-08 | 0.8240 | 0.743 | 0.505 | 0.00121 |
| *Col4a1* | 1.30e-20 | 0.8136 | 0.676 | 0.161 | 1.695e-16 |
| *Col1a1* | 7.01e-10 | 0.8048 | 0.8 | 0.479 | 9.12e-06 |
| *Fhl1* | 5.69e-12 | 0.7733 | 0.59 | 0.208 | 7.41e-08 |
| *Actb* | 1.44e-10 | 0.7712 | 0.867 | 0.568 | 1.87e-06 |
| *Psap* | 9.89e-13 | 0.7357 | 0.905 | 0.745 | 1.29e-08 |
| *Cryab* | 4.70e-12 | 0.7247 | 0.981 | 0.927 | 6.12e-08 |
| *Flnc* | 1.08e-14 | 0.7211 | 0.705 | 0.266 | 1.40e-10 |
| *Eef1a1* | 2.46e-07 | 0.7099 | 0.819 | 0.677 | 0.00320 |
| *Uqcr11* | 2.22e-19 | 0.6736 | 1 | 0.99 | 2.89e-15 |
| *Ctsb* | 3.02e-12 | 0.6639 | 0.819 | 0.443 | 3.93e-08 |
| *Col6a3* | 9.17e-16 | 0.6617 | 0.524 | 0.104 | 1.19e-11 |
| *Col4a2* | 2.12e-14 | 0.6599 | 0.581 | 0.172 | 2.75e-10 |
| *Tmsb4x* | 1.26e-08 | 0.6528 | 0.886 | 0.594 | 0.00016 |
| *Jph2* | 4.11e-11 | 0.6430 | 0.819 | 0.49 | 5.34e-07 |
| *Mustn1* | 5.81e-10 | 0.6389 | 0.59 | 0.25 | 7.55e-06 |
| *Col6a2* | 2.30e-17 | 0.6385 | 0.6 | 0.13 | 2.99e-13 |
| *Gpx1* | 2.25e-11 | 0.6247 | 0.819 | 0.469 | 2.93e-07 |
| *Mybph* | 1.44e-07 | 0.6178 | 0.848 | 0.594 | 0.00187 |
| *Uqcrq* | 3.15e-13 | 0.6145 | 0.99 | 0.979 | 4.10e-09 |
| *Cilp* | 1.78e-17 | 0.6132 | 0.486 | 0.062 | 2.32e-13 |
| *Thbs4* | 1.02e-10 | 0.6128 | 0.457 | 0.125 | 1.32e-06 |
| *Col1a2* | 3.27e-08 | 0.6126 | 0.924 | 0.641 | 0.00043 |
| *Myog* | 3.52e-22 | 0.5986 | 0.419 | 0 | 4.58e-18 |
| *Pdlim3* | 1.62e-09 | 0.5943 | 0.867 | 0.62 | 2.11e-05 |
| *Cd63* | 3.56e-11 | 0.5806 | 0.714 | 0.323 | 4.64e-07 |

| | | | | | |
|---|---|---|---|---|---|
| *Timm13* | 2.92e-13 | 0.5790 | 0.79 | 0.339 | 3.80e-09 |
| *Hspb8* | 2.642e-10 | 0.5773 | 0.771 | 0.453 | 3.44e-06 |
| *Scn1b* | 4.59e-10 | 0.5766 | 0.943 | 0.682 | 5.97e-06 |
| *Myl12a* | 2.23e-09 | 0.5753 | 0.724 | 0.406 | 2.90e-05 |
| *Ifitm3* | 5.223e-13 | 0.5736 | 0.571 | 0.167 | 6.79e-09 |
| *Uqcrc1* | 1.26e-10 | 0.5716 | 0.905 | 0.708 | 1.63e-06 |

**Differentially expressed genes fibrosis.**

*Table S6.* Top 50 fibrosis markers ranked on average log2FC.

| Gene | p value | avg log2FC | pct.1 | pct.2 | Adjusted p value |
|---|---|---|---|---|---|
| *Col1a1* | 1.36e-40 | 3.3450 | 1 | 0.635 | 1.68e-36 |
| *Col1a2* | 1.81e-29 | 2.6475 | 0.977 | 0.81 | 2.23e-25 |
| *Spp1* | 2.34e-12 | 1.9820 | 0.773 | 0.45 | 2.89e-08 |
| *Col3a1* | 1.75e-17 | 1.7884 | 0.977 | 0.74 | 2.16e-13 |
| *Lyz2* | 3.10e-12 | 1.7594 | 0.875 | 0.655 | 3.82e-08 |
| *Sparc* | 8.93e-21 | 1.7143 | 0.943 | 0.685 | 1.10e-16 |
| *Actb* | 9.46e-24 | 1.5735 | 0.977 | 0.86 | 1.17e-19 |
| *Apoe* | 3.04e-14 | 1.5574 | 0.932 | 0.705 | 3.75e-10 |
| *Bgn* | 1.70e-23 | 1.4714 | 0.761 | 0.19 | 2.10e-19 |
| *Ctsb* | 6.83e-22 | 1.4625 | 0.955 | 0.745 | 8.43e-18 |
| *Ftl1* | 1.64e-12 | 1.4318 | 0.909 | 0.715 | 2.02e-08 |
| *Fmod* | 1.12e-08 | 1.3917 | 0.443 | 0.17 | 0.00014 |
| *Fn1* | 1.46e-20 | 1.3814 | 0.795 | 0.3 | 1.80e-16 |
| *Mgp* | 4.50e-17 | 1.3707 | 0.67 | 0.21 | 5.56e-13 |
| *Thbs4* | 1.18e-20 | 1.3187 | 0.693 | 0.165 | 1.46e-16 |
| *Ctsk* | 6.65e-14 | 1.1726 | 0.568 | 0.165 | 8.21e-10 |
| *Ctsl* | 3.98e-11 | 1.1569 | 0.716 | 0.395 | 4.91e-07 |
| *Psap* | 2.40e-14 | 1.1449 | 0.932 | 0.79 | 2.97e-10 |
| *Vim* | 1.39e-14 | 1.1116 | 0.784 | 0.395 | 1.71e-10 |
| *Tmsb10* | 8.58e-14 | 1.1108 | 0.886 | 0.56 | 1.06e-09 |
| *Lgals3* | 2.13e-11 | 1.0919 | 0.75 | 0.41 | 2.63e-07 |
| *H2-D1* | 4.82e-16 | 1.0637 | 0.818 | 0.4 | 5.95e-12 |
| *Acp5* | 3.51e-11 | 1.0383 | 0.375 | 0.07 | 4.33e-07 |
| *Serpinf1* | 2.57e-17 | 1.0343 | 0.682 | 0.195 | 3.17e-13 |
| *Ctsz* | 1.47e-16 | 1.0299 | 0.67 | 0.205 | 1.81e-12 |
| *Grn* | 1.39e-17 | 1.0058 | 0.648 | 0.165 | 1.71e-13 |
| *Cd74* | 1.76e-12 | 0.9903 | 0.682 | 0.28 | 2.17e-08 |
| *Mmp2* | 1.82e-13 | 0.9580 | 0.557 | 0.145 | 2.25e-09 |
| *Crip1* | 5.74e-16 | 0.9443 | 0.784 | 0.33 | 7.08e-12 |
| *Ckb* | 1.22e-14 | 0.9399 | 0.682 | 0.255 | 1.51e-10 |
| *Cxcl14* | 2.17e-17 | 0.9373 | 0.602 | 0.135 | 2.68e-13 |
| *Col6a1* | 8.93e-22 | 0.9343 | 0.739 | 0.17 | 1.10e-17 |
| *S100a6* | 5.87e-14 | 0.9299 | 0.909 | 0.615 | 7.25e-10 |
| *C3* | 3.12e-12 | 0.9190 | 0.511 | 0.145 | 3.85e-08 |
| *Anxa2* | 1.62e-15 | 0.9145 | 0.636 | 0.175 | 1.99e-11 |
| *Postn* | 3.30e-23 | 0.9120 | 0.591 | 0.065 | 4.07e-19 |
| *Ctss* | 4.59e-12 | 0.8956 | 0.557 | 0.175 | 5.66e-08 |
| *Actg1* | 1.76e-14 | 0.8921 | 0.909 | 0.555 | 2.17e-10 |
| *Eef1a1* | 1.03e-12 | 0.8767 | 0.909 | 0.65 | 1.27e-08 |
| *Col6a2* | 1.56e-19 | 0.8760 | 0.682 | 0.155 | 1.93e-15 |
| *Igfbp5* | 2.46e-09 | 0.8723 | 0.67 | 0.325 | 3.04e-05 |

| | | | | | |
|---|---|---|---|---|---|
| *Serpinh1* | 9.84e-18 | 0.8714 | 0.625 | 0.15 | 1.22e-13 |
| *Chad* | 1.71e-11 | 0.8677 | 0.307 | 0.035 | 2.11e-07 |
| *B2m* | 3.68e-16 | 0.8572 | 0.739 | 0.255 | 4.55e-12 |
| *Tmsb4x* | 1.05e-12 | 0.8537 | 0.955 | 0.675 | 1.30e-08 |
| *Col6a3* | 9.35e-15 | 0.8534 | 0.591 | 0.15 | 1.15e-10 |
| *Pcolce* | 5.42e-18 | 0.8458 | 0.659 | 0.175 | 6.69e-14 |
| *Mmp12* | 9.53e-15 | 0.8414 | 0.58 | 0.15 | 1.18e-10 |
| *Timp2* | 1.18e-16 | 0.8396 | 0.705 | 0.22 | 1.43e-12 |
| *S100a4* | 2.67e-16 | 0.8315 | 0.568 | 0.12 | 3.30e-12 |

#### Differentially expressed genes calcification.

*Table S7.* Top 50 calcification markers ranked on average log2FC.

| Gene | p value | avg log2FC | pct.1 | pct.2 | Adjusted p value |
|---|---|---|---|---|---|
| *Spp1* | 4.68e-54 | 2.6642 | 0.9 | 0.475 | 5.78e-50 |
| *Lyz2* | 2.47e-54 | 1.9674 | 0.98 | 0.688 | 3.05e-50 |
| *Actb* | 1.88e-67 | 1.8507 | 0.99 | 0.833 | 2.32e-63 |
| *Ctsb* | 3.97e-70 | 1.7721 | 1 | 0.748 | 4.91e-66 |
| *Ftl1* | 1.66e-38 | 1.6713 | 0.965 | 0.743 | 2.05e-34 |
| *Col3a1* | 1.91e-32 | 1.6164 | 0.945 | 0.787 | 2.35e-28 |
| *Apoe* | 3.77e-42 | 1.6116 | 0.95 | 0.716 | 4.65e-38 |
| *Psap* | 5.24e-59 | 1.5832 | 0.99 | 0.784 | 6.47e-55 |
| *Col1a1* | 9.13e-49 | 1.5433 | 0.98 | 0.862 | 1.13e-44 |
| *Tmsb10* | 4.09e-53 | 1.5138 | 0.965 | 0.615 | 5.05e-49 |
| *Mgp* | 2.11e-111 | 1.4931 | 0.865 | 0 | 2.60e-107 |
| *Lgals3* | 7.77e-46 | 1.4522 | 0.855 | 0.429 | 9.60e-42 |
| *Col1a2* | 1.48e-37 | 1.4497 | 0.975 | 0.844 | 1.83e-33 |
| *Fn1* | 7.52e-50 | 1.4274 | 0.865 | 0.344 | 9.29e-46 |
| *Ckb* | 1.35e-48 | 1.4210 | 0.835 | 0.328 | 1.66e-44 |
| *Bgn* | 1.17e-42 | 1.3942 | 0.765 | 0.275 | 1.44e-38 |
| *Ctsl* | 3.52e-37 | 1.3655 | 0.855 | 0.427 | 4.34e-33 |
| *Mmp12* | 1.97e-40 | 1.3498 | 0.66 | 0.177 | 2.44e-36 |
| *Ctsz* | 2.00e-49 | 1.3035 | 0.83 | 0.31 | 2.48e-45 |
| *Ctsk* | 1.49e-33 | 1.2342 | 0.65 | 0.209 | 1.84e-29 |
| *H2-D1* | 1.18e-43 | 1.2177 | 0.88 | 0.436 | 1.46e-39 |
| *Ctss* | 1.45e-41 | 1.1599 | 0.725 | 0.22 | 1.79e-37 |
| *Vim* | 6.95e-35 | 1.1357 | 0.855 | 0.452 | 8.59e-31 |
| *Sparc* | 6.50e-26 | 1.1219 | 0.935 | 0.757 | 8.03e-22 |
| *Crip1* | 4.27e-43 | 1.1203 | 0.88 | 0.408 | 5.27e-39 |
| *Anxa2* | 1.094e-39 | 1.1067 | 0.735 | 0.239 | 1.35e-35 |
| *Grn* | 8.38e-46 | 1.0836 | 0.78 | 0.234 | 1.03e-41 |
| *Actg1* | 3.13e-41 | 1.0757 | 0.945 | 0.603 | 3.86e-37 |
| *S100a6* | 1.17e-39 | 1.0685 | 0.93 | 0.633 | 1.44e-35 |
| *Cxcl14* | 1.80e-41 | 1.0604 | 0.7 | 0.183 | 2.23e-37 |
| *Cd74* | 1.065e-32 | 1.0424 | 0.755 | 0.321 | 1.31e-28 |
| *Acp5* | 6.81e-23 | 1.0376 | 0.44 | 0.112 | 8.41e-19 |
| *Tmsb4x* | 1.15e-35 | 1.0372 | 0.955 | 0.711 | 1.42e-31 |
| *Ctsd* | 1.10e-31 | 1.0349 | 0.955 | 0.709 | 1.36e-27 |
| *Gpx1* | 2.11e-33 | 1.0244 | 0.915 | 0.589 | 2.60e-29 |
| *Atp6v0c* | 6.27e-35 | 0.9659 | 0.835 | 0.42 | 7.74e-31 |
| *Fabp5* | 2.75e-30 | 0.9655 | 0.675 | 0.25 | 3.39e-26 |
| *Postn* | 2.78e-42 | 0.9523 | 0.645 | 0.138 | 3.44e-38 |
| *Eef1a1* | 2.85e-30 | 0.9460 | 0.925 | 0.679 | 3.52e-26 |

| | | | | | |
|---|---|---|---|---|---|
| *B2m* | 2.59e-37 | 0.9261 | 0.795 | 0.314 | 3.20e-33 |
| *Ptma* | 2.51e-38 | 0.9168 | 0.825 | 0.339 | 3.10e-34 |
| *Cd63* | 2.28e-33 | 0.9009 | 0.835 | 0.413 | 2.74e-29 |
| *C3* | 3.41e-25 | 0.8907 | 0.555 | 0.179 | 4.22e-21 |
| *Cyba* | 1.66e-40 | 0.8846 | 0.67 | 0.161 | 2.04e-36 |
| *Myl6* | 6.37e-33 | 0.8798 | 0.88 | 0.491 | 7.87e-29 |
| *Eln* | 2.09e-24 | 0.8715 | 0.415 | 0.087 | 2.58e-20 |
| *C1qa* | 1.01e-39 | 0.8684 | 0.68 | 0.177 | 1.25e-35 |
| *Bc1* | 6.55e-08 | 0.8679 | 0.74 | 0.768 | 0.00081 |
| *Cfl1* | 3.83e-31 | 0.8317 | 0.735 | 0.303 | 4.73e-27 |
| *H2-K1* | 2.46e-41 | 0.8314 | 0.67 | 0.161 | 3.04e-37 |

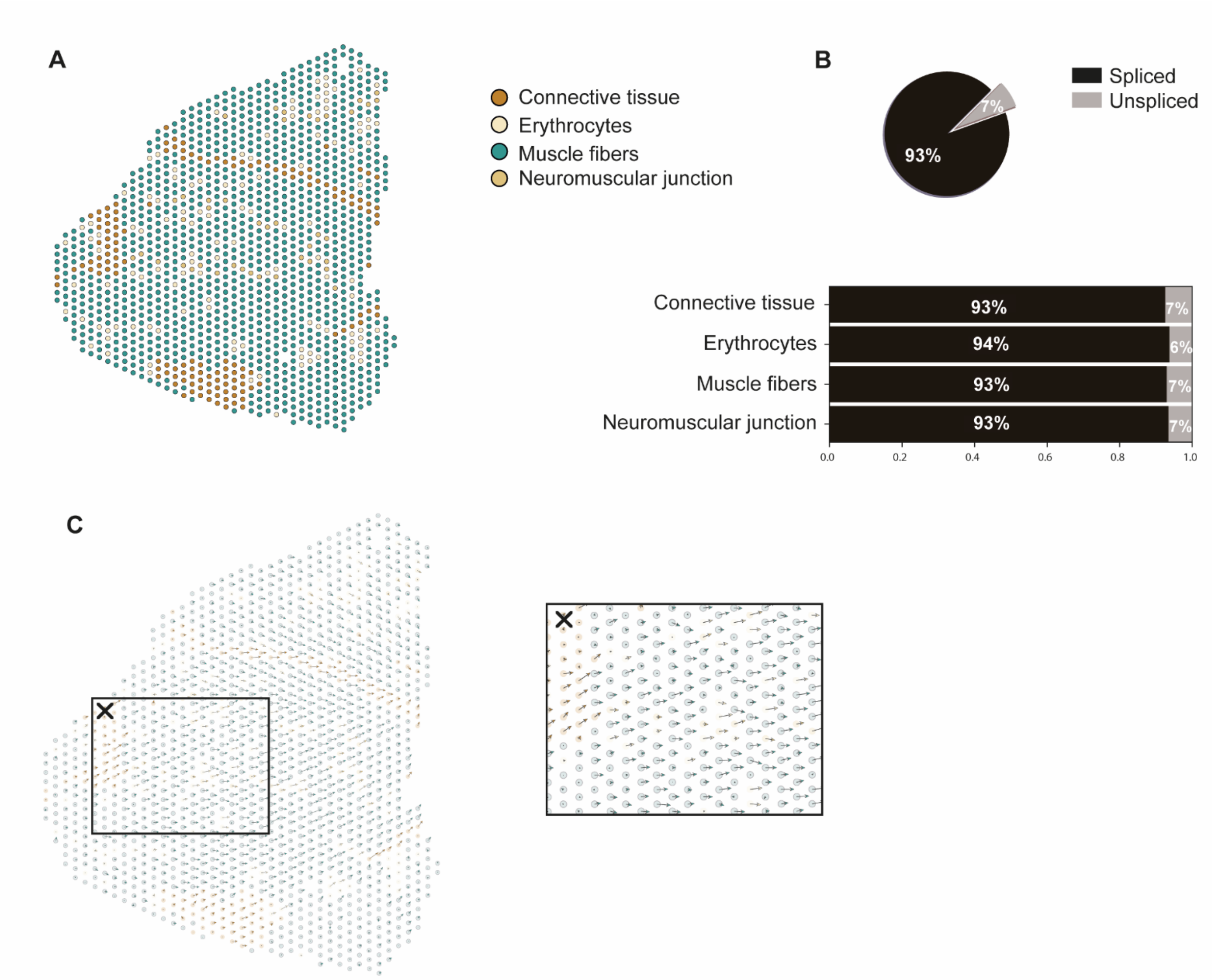


*Figure S6.* **RNA velocity applied on DBA/2J muscle shows that there are no differentiation dynamics present.** *(A) Annotated clusters of DBA/2J as described before (B) Proportion of spliced/unspliced counts in the D2-mdx sample and in its annotated clusters (C) Spatial spot-level RNA velocity vectors showing no differentiation pattern as arrows are indicating the direction and strength of the change in transcriptional state in each spot (✕ box for zoomed-in view).*
